# R-loop homeostasis and cancer mutagenesis promoted by the DNA cytosine deaminase APOBEC3B

**DOI:** 10.1101/2021.08.30.458235

**Authors:** Jennifer L. McCann, Agnese Cristini, Emily K. Law, Seo Yun Lee, Michael Tellier, Michael A. Carpenter, Chiara Beghè, Jae Jin Kim, Matthew C. Jarvis, Bojana Stefanovska, Nuri A. Temiz, Erik N. Bergstrom, Daniel J. Salamango, Margaret R. Brown, Shona Murphy, Ludmil B. Alexandrov, Kyle M. Miller, Natalia Gromak, Reuben S. Harris

**Affiliations:** Howard Hughes Medical Institute, University of Minnesota, Minneapolis, Minnesota, USA, 55455; Masonic Cancer Center, University of Minnesota, Minneapolis, Minnesota, USA, 55455; Institute for Molecular Virology, University of Minnesota, Minneapolis, Minnesota, USA, 55455; Department of Biochemistry, Molecular Biology and Biophysics, University of Minnesota, Minneapolis, Minnesota, USA, 55455; Sir William Dunn School of Pathology, University of Oxford, South Parks Road, Oxford, UK, OX1 3RE; Department of Molecular Biosciences, University of Texas at Austin, Austin, Texas, USA, 78712; Institute for Health Informatics, University of Minnesota, Minneapolis, MN, USA, 55455; Department of Cellular and Molecular Medicine, UC San Diego, La Jolla, California, USA, 92093; Department of Bioengineering, UC San Diego, La Jolla, California, USA, 92093; Moores Cancer Center, UC San Diego, La Jolla, CA, 92037, USA; Livestrong Cancer Institutes, Dell Medical School, University of Texas at Austin, Austin, Texas, USA, 78712

**Author notes:** Equal primary contributions (Jennifer L. McCann, Agnese Cristini). Equal secondary contributions (Emily K. Law, Seo Yun Lee).

**Keywords:** APOBEC mutation signature, APOBEC3B, Cancer mutagenesis, DNA damage, DNA deamination, *Kataegis*, R-loop homeostasis, R-loop mutagenesis, RNA/DNA hybrids

## Abstract

The single-stranded DNA cytosine-to-uracil deaminase APOBEC3B is an antiviral protein implicated in cancer. However, its substrates in cells are not fully delineated. Here, APOBEC3B proteomics reveal interactions with a surprising number of R-loop factors. Biochemical experiments show APOBEC3B binding to R-loops in human cells and *in vitro*. Genetic experiments demonstrate R-loop increases in cells lacking APOBEC3B and decreases in cells overexpressing APOBEC3B. Genome-wide analyses show major changes in the overall landscape of physiological and stimulus-induced R-loops with thousands of differentially altered regions as well as binding of APOBEC3B to many of these sites. APOBEC3 mutagenesis impacts overexpressed genes and splice factor mutant tumors preferentially, and APOBEC3-attributed *kataegis* are enriched in RTCW consistent with APOBEC3B deamination. Taken together with the fact that APOBEC3B binds single-stranded DNA and RNA and preferentially deaminates DNA, these results support a mechanism in which APOBEC3B mediates R-loop homeostasis and contributes to R-loop mutagenesis in cancer.

**Highlights:**

- Unbiased proteomics link antiviral APOBEC3B to R-loop regulation
- Systematic alterations of APOBEC3B levels trigger corresponding changes in R-loops
- APOBEC3B binds R-loops in living cells and *in vitro*
- Bioinformatics analyses support an R-loop deamination and mutation model

## Introduction

The APOBEC3 family of single-stranded (ss)DNA cytosine deaminases are critical players in the overall innate immune response to viral infection (reviewed by refs.^1, 2^). Popularized initially by potent HIV-1 restriction activity^3–6^, the seven human APOBEC3 enzymes collectively elicit activity against a broad number of DNA-based viruses including retroviruses (HIV-1, HTLV-1), hepadnaviruses (HBV), papillomaviruses (HPV), parvoviruses (AAV), polyomaviruses (JC and BKPyV), and herpesviruses (EBV, HSV-1) (reviewed by refs.^1,2^). An important biochemical feature of this family of enzymes is an intrinsic preference for different nucleobases immediately 5’ of target cytosines^7–11^. For example, APOBEC3B (A3B) and APOBEC3A (A3A) preferentially deaminate cytosines in 5’TC motifs and the related antibody gene diversification enzyme AID targets 5’AC and 5’GC motifs.

In addition to beneficial functions in innate and adaptive immunity, multiple DNA cytosine deaminases have detrimental roles in cancer mutagenesis (reviewed by refs.^1,12–17^). Misprocessing of AID-catalyzed deamination events in antibody gene variable and switch regions can result in DNA breaks and chromosomal translocations in B cell malignancies (reviewed by ref.^17^). Off-target deamination of other genes also occurs at lower frequencies and the resulting mutations may contribute to B cell lineage-derived leukemias and lymphomas^18,19^. In comparison, large-scale tumor sequencing projects have reported an APOBEC mutation signature in a wide variety of cancer types including those derived from breast, lung, head/neck, cervix, and bladder tissues (see recent pan-cancer analysis^20^ and reviews above). In cancer, the APOBEC mutation signature is defined as C-to-T transitions and C-to-G transversions in 5’TCA and 5’TCT motifs to help distinguish from other mutational processes^21–25^. Recent estimates indicate that APOBEC enzymes are the second most prevalent mutation-generating process in human cancer following clock-like mutagenesis which is attributed to ageing^20^.

Despite extensive documentation of the APOBEC mutation signature in cancer, the precise molecular mechanisms governing this mutational process are still unclear. One challenge is the possibility that at least two enzymes, A3B and A3A, combine in different ways to generate the overall signature (*e.g.*, refs.^21–23,26–28^). However, a few mechanistic insights have still been inferred from the physical characteristics of genomes with, for instance, APOBEC mutations associating with chromosomal DNA replication and specifically with C-to-U deamination events in the lagging strand template^29–34^. Other genomic features and processes with exposed ssDNA may be similarly prone to APOBEC mutagenesis such as single-stranded loop regions of DNA hairpins^35,36^ and ssDNA tracts in recombination and repair reactions, which sometimes manifest clusters of strand-coordinated APOBEC3-attributed mutations (*aka. kataegis*; *e.g.*, refs.^25,35,37^). Together, these studies have indicated a passive diffusion mechanism in which simple expression of A3B and/or A3A leads to random encounters with any exposed ssDNA followed in some instances by processive local deamination.

Another potential substrate for APOBEC enzymes is ssDNA exposed by R-loop formation. R-loops occur when nascent RNA re-anneals to the transcribed DNA strand, creating a three-stranded structure containing an RNA/DNA hybrid and a displaced non-transcribed ssDNA strand (reviewed by refs.^38–40^). R-loops are critical for the specialized mechanism of AID-catalyzed antibody gene hypermutation and class switch recombination in B lymphocytes (reviewed by refs.^41,42^). R-loops also represent a prominent source of genome instability associated with human disease, including cancer (*e.g.*, refs.^43–45^; reviewed by refs.^39,40,46–51^). However, evidence linking APOBEC enzymes to R-loop-associated mutation and genome instability is presently lacking. A potential clue came from recent studies showing a mismatch repair-dependent synthetic lethal interaction between A3B activity and uracil excision repair disruption^52^. This report suggested that the toxic lesions may represent U/G mispairs, which are unlikely to arise through replication but may be created by transcription-associated processes, such as C-to-U deamination within a R-loop.

A3B is strongly implicated in cancer mutagenesis based on several criteria including nuclear localization, overexpression in tumors, upregulation by cancer-causing viruses such as HPV, and associations with clinical outcomes (reviewed by refs.^1,12–14^). To gain deeper insights into the pathological role of A3B in cancer mutagenesis, an unbiased affinity purification and mass spectrometry (AP-MS) approach was used to identify A3B-interacting proteins. Two dozen proteins were recovered in biologically-independent experiments, and 60% of the resulting high-confidence interactors had been reported previously as R-loop-associated factors in RNA/DNA hybrid AP-MS experiments^53^. A comprehensive series of genetic, cell biology, biochemistry, genomic, and bioinformatic studies was therefore undertaken to investigate the relationship between A3B and R-loops. Surprisingly, the results indicated that A3B functions in R-loop homeostasis. Moreover, R-loop regions impacted by cellular A3B show significant enrichments for APOBEC3 signature mutations including *kataegis* in breast tumors. Altogether, these results reveal an unanticipated role for A3B in R-loop biology and a distinct mechanism of transcription-associated cancer mutagenesis.

## Results

### APOBEC3B interacts with R-loops and R-loop-associated proteins

Little is known about the molecular processes that serve to regulate the mutagenic activities of A3B. To address this gap in knowledge, a functional A3B-2xStrep-3xFlag construct (hereafter A3B-SF) was expressed in 293T cells, anti-Strep affinity-purified, and subjected to mass spectrometry to identify interacting proteins (AP-MS workflow schematic in **Fig. S1a**). This procedure included exogenous RNase A and high salt concentrations to enrich for direct and strong interactions, respectively. Immunoblots, Coomassie gels, and DNA deaminase activity assays were used to validate the presence, enrichment, and activity of affinity-purified A3B (**Fig. S1b-d**). Expression of an eGFP-SF construct and an empty 2xStrep-3xFlag vector was analyzed in parallel as negative controls.

Six independent AP-MS experiments yielded a total of 24 specific A3B-interacting proteins (**Table S1**; independent confirmation in **Fig. S1e-f**). These proteins were abundant in all 6 A3B-SF data sets and, importantly, absent in GFP-SF or empty vector datasets. Only CDK4, which regulates A3B localization during the cell cycle^54^, and hnRNPUL1, which tethered to Cas9 is capable of targeting A3B for base editing^55^, had been reported previously. However, literature analyses revealed that 60% of these A3B interactors had been found independently in S9.6 AP-MS experiments^53^ (**Fig. 1a-b**). As the S9.6 mAb binds strongly to RNA/DNA hybrid structures^56,57^, this significant interactome overlap suggested that A3B may also interact with R-loops. To test this hypothesis, interactions between A3B and multiple R-loop factors were confirmed in co-immunoprecipitation (co-IP) experiments. For example, doxycycline (Dox)-inducible A3B-eGFP was immunoprecipitated from MCF10A cells with an anti-eGFP antibody and the R-loop-associated protein hnRNPUL1 was detected by immunoblotting (system validation in **Fig. 1c** and representative co-IP in **Fig. 1d**). Parallel slot blots showed that R-loops also co-purified with A3B-eGFP, and the RNase H sensitivity of the signal demonstrated the specific detection of RNA/DNA hybrids (**Fig. 1d**).

**Fig. 1.**
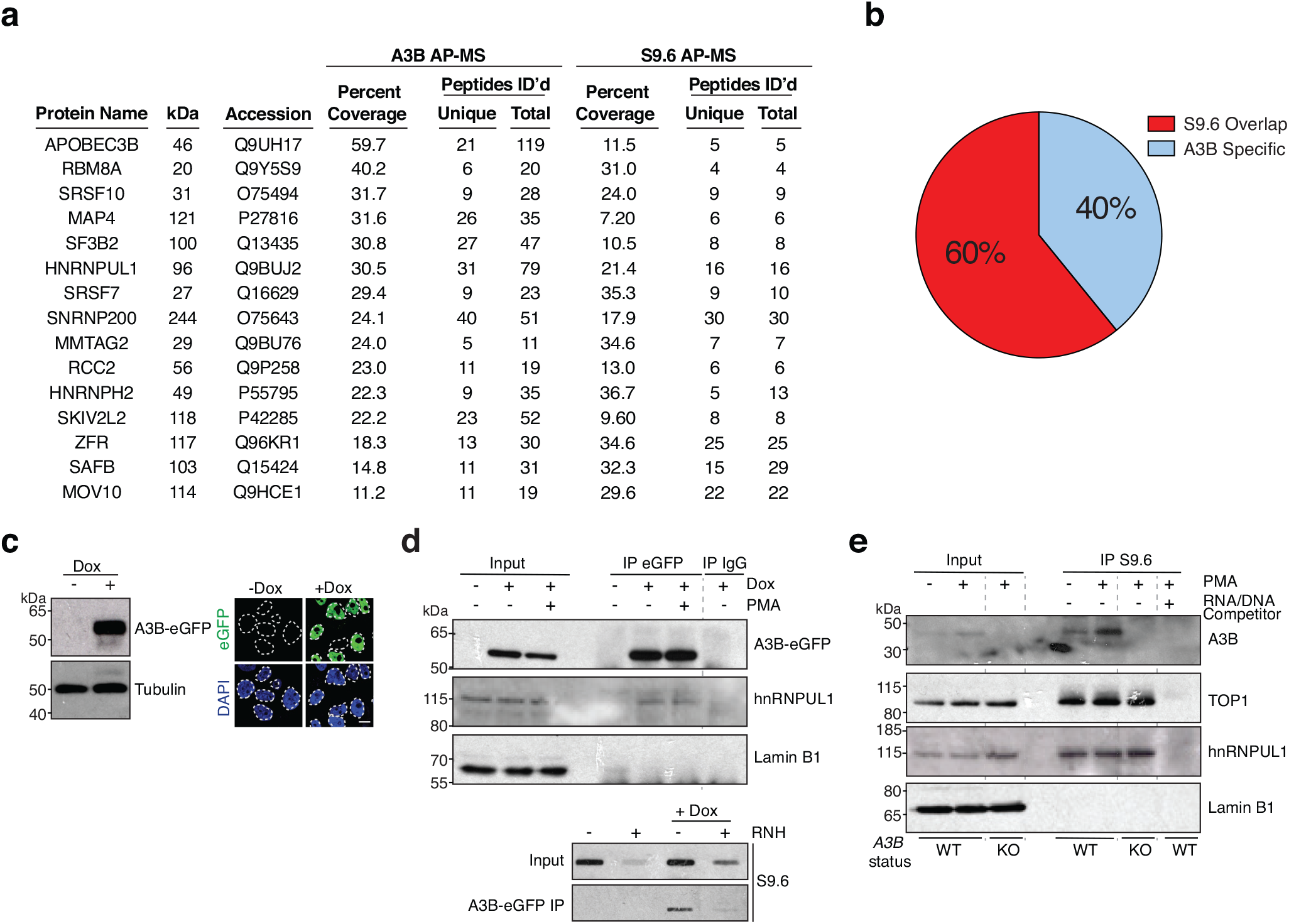
A3B interacts with R-loop-associated proteins. **a**, Information on shared proteins between A3B and S9.6 AP-MS datasets. **b**, Pie chart depicting overlap between A3B and S9.6 AP-MS datasets. **c**, Immunoblot and IF microscopy analysis of MCF10A-TREx-A3B-eGFP cells treated with vehicle or Dox (1 µg/mL, 24 hrs). Nuclear A3B-eGFP is green and nuclear DNA is blue (DAPI; representative images; 10 μm scale bar). **d**, Immunoblots of A3B-eGFP or IgG IP from TREx-A3B-eGFP MCF10A cells +/- Dox (1 μg/ml, 24 hrs), treated with PMA (25 ng/ml, 2 hrs), and probed with indicated antibodies (upper images). Slot blot of A3B-eGFP IP from TREx-A3B-eGFP MCF10A cells +/- Dox (1 μg/ml, 24 hrs) probed with S9.6 antibody (lower images). RNase H (RNH) treatment confirmed specificity of R-loop signal. **e**, Immunoblots of the indicated proteins in input lysates and S9.6 IP reactions from MCF10A (WT) or *A3B*-null derivatives (KO) treated with PMA (25 ng/ml, 5 hrs). R-loop specificity is indicated by signal depletion by RNA/DNA hybrid competitor.

The S9.6 mAb was then used to IP RNA/DNA hybrids from MCF10A cells treated with phorbol 12-myristate 13-acetate (PMA) to induce endogenous *A3B* expression^58^. Immunoblotting confirmed enrichment of an established R-loop interacting protein, TOP1^59,60^, and a shared R-loop and A3B interactor, hnRNPUL1, in all S9.6 IP reactions except those saturated with a synthetic RNA/DNA hybrid competitor (**Fig. 1e**). The non-R-loop-interacting protein Lamin B1 served as a negative control for these IP reactions. Endogenous A3B copurified with R-loop structures in basal non-induced conditions and this interaction increased following PMA treatment (**Fig. 1e**). Importantly, no A3B signal was detected in S9.6 pull downs from *A3B*-null MCF10A cells, which demonstrated interaction specificity (**Fig. 1e**; *A3B* knockout construction and validation in **Fig. S2a-d**).

### Elevated nuclear R-loop levels in *A3B*-null and *A3B*-depleted cells

To investigate a potential role for A3B in R-loop biology, R-loop levels were quantified in MCF10A and its *A3B*-null derivative. First, nucleoplasmic S9.6 staining intensity was measured by immunofluorescence (IF) confocal microscopy. These experiments revealed a strong increase in nucleoplasmic S9.6 fluorescence in *A3B*-null cells in comparison to parental MCF10A cells (**Fig. 2a-b**). Second, S9.6 dot blot experiments confirmed elevated R-loop levels in the *A3B*-null MCF10A cells in comparison to the parental line (**Fig. 2c-d**). Importantly, in both series of experiments, RNase H treatment eliminated the increase in R-loop signals observed in the absence of endogenous A3B. S9.6 signal detecting nucleolar rRNA is mostly insensitive to RNase H treatment, as reported^61^, and is also largely unaffected by A3B.

**Fig. 2.**
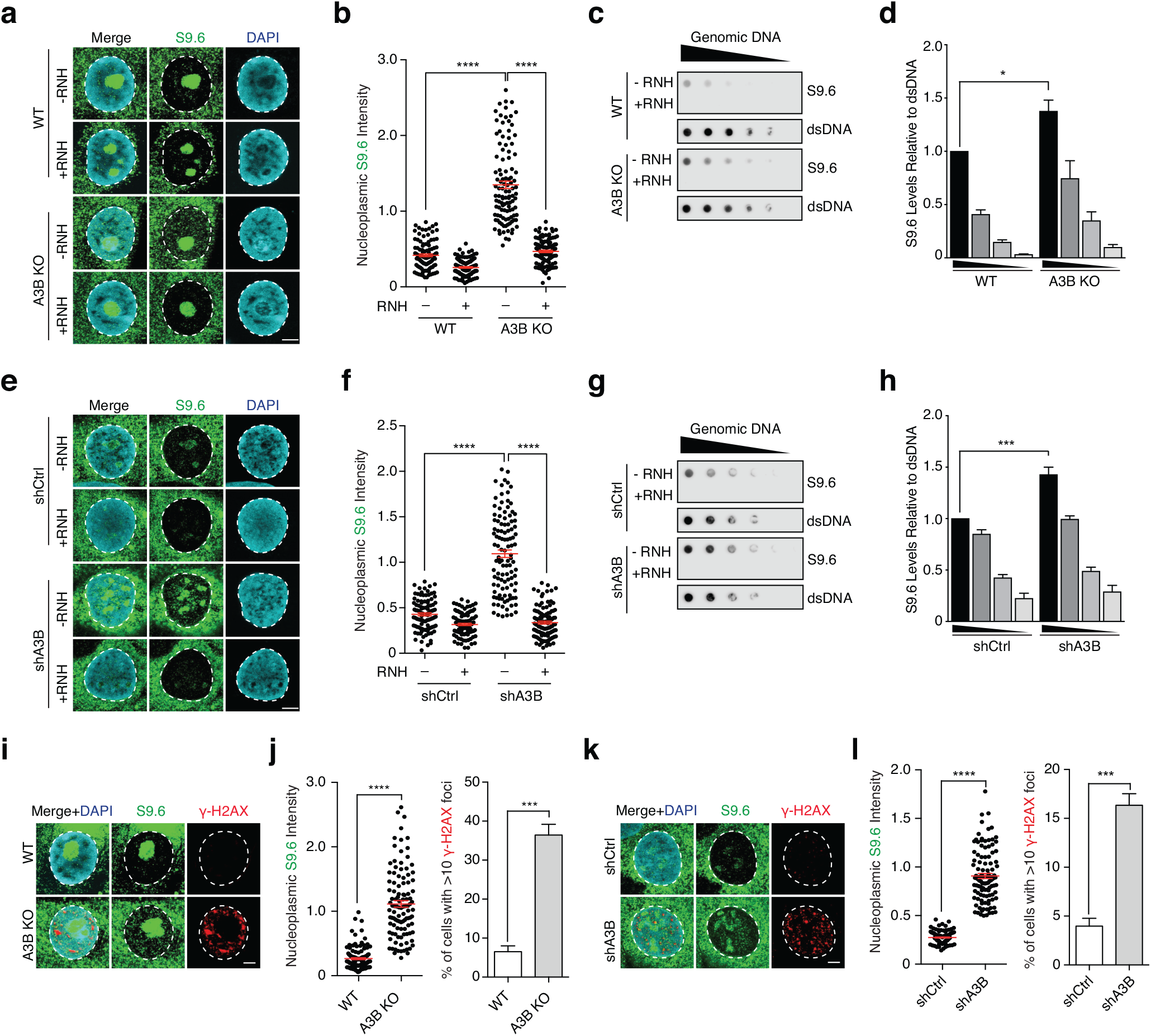
Elevated nuclear R-loop levels in *A3B*-null and *A3B*-depleted cells. **a-b**, IF images and quantification of MCF10A (WT) and *A3B*-null (KO) cells stained with S9.6 (green) and DAPI (blue). Diminution of nucleoplasmic S9.6 signal by exogenous RNase H (RNH) confirmed signal specificity (representative images; 5 μm scale bar; n > 100 nuclei per condition; red bars represent mean ± SEM; ****, *P* < 0.0001 by Mann-Whitney test). **c-d**, S9.6 dot blot analysis of a MCF10A (WT) and *A3B*-null (KO) genomic DNA dilution series +/- exogenous RNase H (RNH; representative images). Parallel dsDNA dot blots provided a loading control. Quantification normalized to the most concentrated WT signal (n ≥ 4; mean ± SEM; *, *P* < 0.05 by two-tailed unpaired t-test). **e-f**, IF images and quantification of U2OS shCtrl and shA3B cells stained with S9.6 (green) and DAPI (blue). Diminution of nucleoplasmic S9.6 signal by exogenous RNase H (RNH) confirmed signal specificity (representative images; 5 μm scale bar; n > 100 nuclei per condition; red bars represent mean ± SEM; ****, *P* < 0.0001 by Mann-Whitney test). **g-h**, S9.6 dot blot analysis of a U2OS shCtrl and shA3B genomic DNA dilution series +/- exogenous RNase H (RNH; representative images). Parallel dsDNA dot blots provided a loading control. Quantification normalized to the most concentrated shCtrl signal (n ≥ 4; mean ± SEM; ***, *P* < 0.001 by two-tailed unpaired t-test). **i-j**, IF images and quantification of MCF10A (WT) and *A3B*-null (KO) cells stained with S9.6 (green), DAPI (blue), and γ-H2AX (representative images; 5 μm scale bar; n > 100 nuclei per condition; red bars represent mean ± SEM; ****, *P* < 0.0001 by Mann-Whitney test, ***, *P* < 0.001 by two-tailed unpaired t-test). **k-l**, IF images and quantification of U2OS shCtrl and shA3B cells stained with S9.6 (green), DAPI (blue), and γ-H2AX (representative images; 5 μm scale bar; n > 100 nuclei per condition; red bars represent mean ± SEM; ****, *P* < 0.0001 by Mann-Whitney test, ***, *P* < 0.001 by two-tailed unpaired t-test).

To address whether the relationship between A3B and R-loop levels may be a general mechanism, analogous S9.6 IF microscopy and dot blot experiments were done using U2OS cells, which express high levels of endogenous A3B^54,62,63^. U2OS cells were transduced with a validated shRNA to stably deplete endogenous *A3B* or a non-targeting control shRNA^22,64^ (*A3B* knockdown validation in **Fig. S2e-g**). In these experiments, *A3B* knockdown caused a strong increase in nucleoplasmic S9.6 staining intensity by IF microscopy in comparison to control transduced cells analyzed in parallel (**Fig. 2e-f**). As above, nearly all nucleoplasmic S9.6 staining was ablated by RNase H treatment, indicating that the signal is specific to RNA/DNA hybrids. An increase in RNA/DNA hybrid signal was also obtained in S9.6 dot blot experiments from *A3B*-depleted versus control transduced cells and, again, specificity was confirmed by RNase H treatment (**Fig. 2g-h**). R-loop imbalances are a known source of DNA damage^38–40^, and elevated R-loop levels in *A3B*-null MCF10A and *A3B*-depleted U2OS cells triggered concomitant increases in DNA break formation as evidenced by γ-H2AX staining (**Fig. 2i-l**). However, it is important to note that these elevated levels of R-loops and DNA damage did not alter overall rates of DNA replication (EdU incorporation) or cell cycle progression (**Fig. S2h-k**).

### A3B reduces nuclear R-loop levels through a deamination-dependent mechanism

Given results above showing increased R-loop accumulation upon *A3B* loss, we next asked whether A3B overexpression might have the opposite effect and suppress R-loop formation. U2OS cells were transfected with an expression vector for A3B-eGFP or eGFP as a negative control, incubated 24 hrs to allow for protein expression, treated 4 hrs with the Bromodomain and Extra-Terminal (BET) protein family inhibitor JQ1 to enhance R-loop formation^44,65,66^, and then analyzed by IF confocal microscopy for S9.6 staining. In comparison to the eGFP control, A3B-eGFP expression caused a significant decrease in the average intensity of nucleoplasmic S9.6 levels (**Fig. 3a-b**). Interestingly, expression of A3A, which is more active than A3B in ssDNA deamination assays^67,68^, had no effect suggesting a specific R-loop role for A3B (**Fig. 3a-b**).

**Fig. 3.**
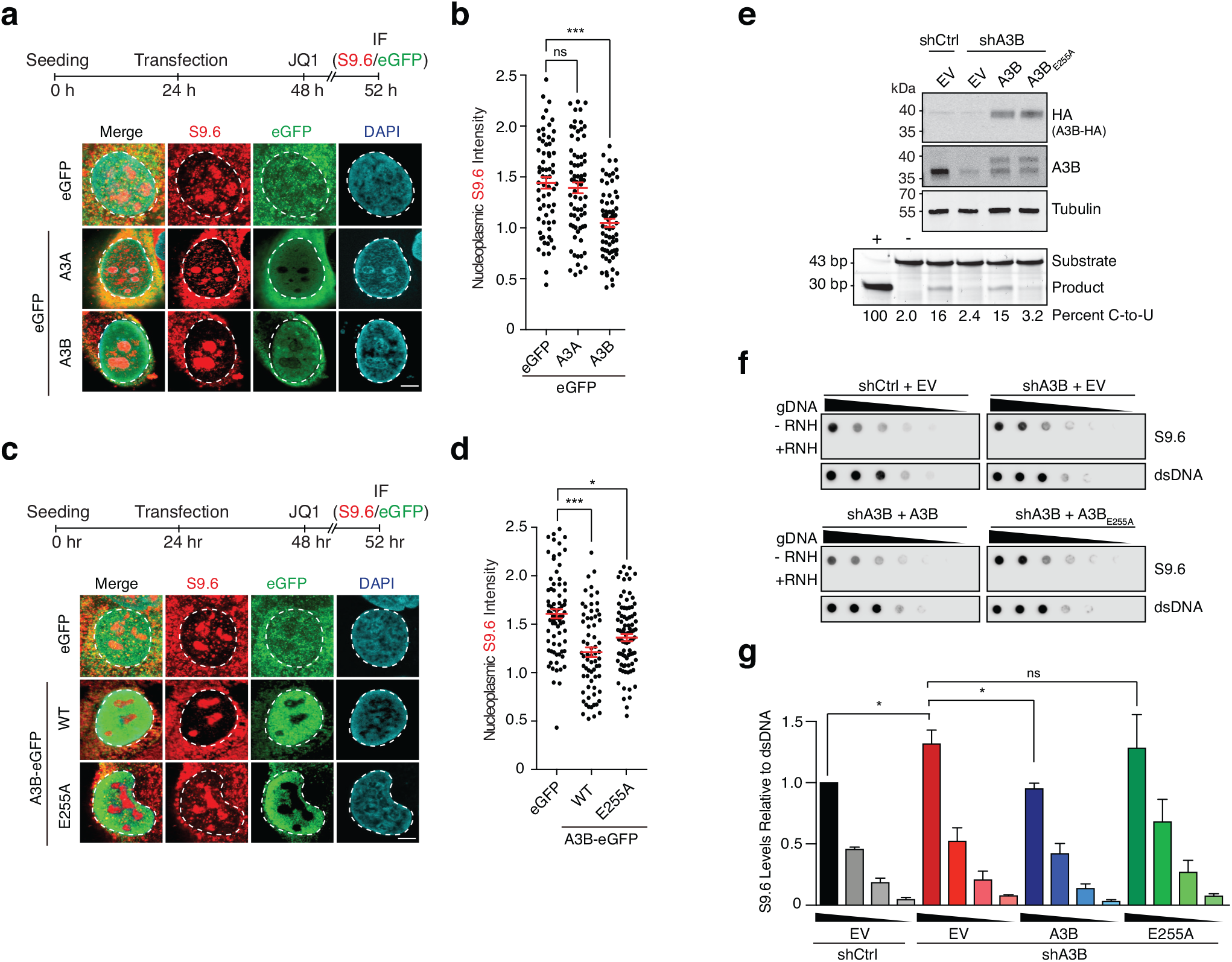
A3B overexpression reduces nuclear R-loop levels. **a-d**, IF images (a and c) and quantification (b and d) of U2OS cells expressing the denoted eGFP construct (green) and stained with S9.6 (red) and DAPI (blue). Upper panel shows experimental workflow and lower panel representative images (5 μm scale bar; n > 60 nuclei per condition; red bars represent mean ± SEM; *, *P* < 0.05 and ***, *P* < 0.001 by one-way ANOVA; ns, not significant). **e**, Immunoblots of U2OS shCtrl or shA3B cells complemented with empty vector (EV), A3B-HA, or A3B-E255A-HA. The lower image shows the results of a DNA deaminase activity assay with extracts from the indicated cell lines (reaction quantification below with purified A3A as a positive control and reaction buffer as a negative control). **f-g**, Dot blot analysis of U2OS shCtrl or shA3B cells complemented with empty vector (EV), A3B-HA, or A3B-E255A-HA. A genomic DNA dilution series +/- exogenous RNase H (RNH) was probed with either S9.6 antibody or dsDNA antibody as a loading control (representative images). Quantification normalized to the most concentrated shCtrl signal (n = 3; mean ± SEM; *, *P* < 0.05 by two-tailed unpaired t-test; ns, not significant).

The hallmark biochemical activity of A3B is ssDNA C-to-U deamination (*e.g.*, refs.^22,67,68^; reviewed by refs.^13,69,70^). To determine whether deaminase activity is required for reducing nuclear R-loop levels, U2OS cells were transfected with constructs expressing A3B-eGFP or a single amino acid substitution catalytic mutant E255A, which prevents A3B from deprotonating water and creating the hydroxide ion required for C-to-U deamination (reviewed by refs.^13,69,70^). Importantly, wild-type A3B caused a significant reduction in nucleoplasmic R-loop levels as quantified by S9.6 staining and IF microscopy, whereas the catalytic mutant had a less pronounced effect despite similar expression levels (**Fig. 3c-d** and data not shown). However, because U2OS cells already express high levels of endogenous A3B (above) and APOBEC3 enzymes including A3B are reported to oligomerize^68,71–75^, this intermediate phenotype may be due to oligomerization between overexpressed mutant and endogenous A3B.

Therefore, a series of genetic complementation experiments was performed to compare the activities of wild-type A3B and the E255A catalytic mutant in cells lacking endogenous A3B. First, endogenous *A3B* was depleted from U2OS cells as above, which resulted in lower A3B protein and activity levels (**Fig. 3e**). Second, *A3B*-depleted U2OS cells were stably transfected with shRNA-resistant constructs expressing HA-tagged wild-type A3B, A3B-E255A, or an empty vector control (**Fig. 3e**). The wild-type A3B enzyme but not the E255A catalytic mutant restored ssDNA deaminase activity as expected (**Fig. 3e**). Third, R-loop levels were analyzed by S9.6 dot blot assays. Importantly, complementation with wild-type A3B rescued the effect of *A3B* depletion and caused a significant reduction in R-loops (**Fig. 3f-g**). In contrast, cells complemented with similar levels of the catalytic mutant protein, A3B-E255A, showed no significant change in R-loop levels demonstrating a deamination-dependent mechanism.

### APOBEC3B-regulated R-loops are transcription-dependent

R-loop formation normally occurs during the process of transcription when a nascent RNA hybridizes with the template DNA strand and displaces the non-template ssDNA behind the elongating RNA polymerase (reviewed by refs.^38,39,76^). To confirm that A3B-associated R-loops are transcription-dependent, shCtrl and shA3B U2OS cells were treated with the global transcription inhibitor triptolide^77,78^ and IF microscopy analyses were used to quantify R-loop levels. As above, *A3B* depletion resulted in an increase in nucleoplasmic R-loop levels, and triptolide abolished this effect (**Fig. 4a-b**). Dot blot analysis of RNA/DNA hybrids yielded similar results (**Fig. 4c-d**). Complementary data were obtained using a different transcription inhibitor, flavopiridol, which specifically inhibits the CDK9 kinase subunit of the positive transcription elongation factor b (P-TEFb) of RNA polymerase II (*i.e.*, flavopiridol inhibits CDK9, blocks transcription elongation, and concomitantly blocks R-loop accumulation^79^) (**Fig. 4c-d**). These results confirmed that A3B-targeted R-loops are indeed transcription-dependent.

**Fig. 4.**
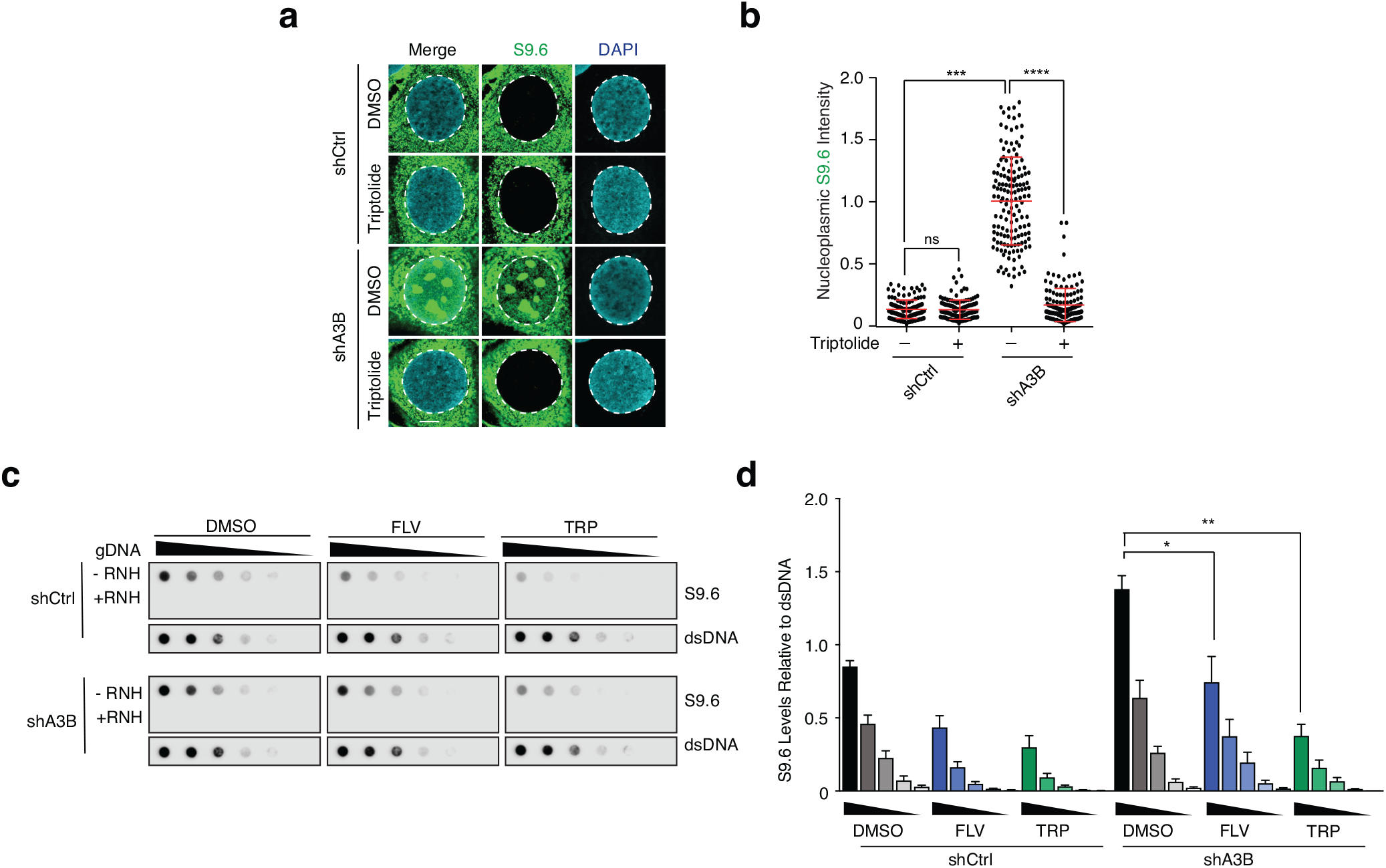
A3B-regulated R-loops are transcription-dependent. **a-b**, IF images and quantification of U2OS shCtrl and shA3B cells treated with triptolide (1 μM, 4 hrs) and subsequently stained with S9.6 (green) and DAPI (blue; representative images; 5 μm scale bar; n > 130 nuclei per condition; red bars represent mean ± SEM; ***, *P* < 0.001 and ****, *P* < 0.0001 by one-way ANOVA; ns, not significant). **c-d**, S9.6 dot blot analysis of a genomic DNA dilution series +/- RNH from U2OS shCtrl or shA3B cells treated with triptolide (TRP; 1 μM, 4 hrs) or flavopiridol (FLV; 1 μM, 1 hr; representative images). Parallel dsDNA dot blots provided a loading control. Quantification was normalized to the most concentrated shCtrl/DMSO signal (n = 3; mean ± SEM; *, *P* < 0.05; **, *P* < 0.01 by two-tailed unpaired t-test).

### APOBEC3B alters the genome-wide distribution of R-loops

The aforementioned IF confocal microscopy and dot blot results suggested a role for A3B in regulating R-loop levels genome-wide. This possibility was investigated using DNA/RNA hybrid IP (DRIP)-seq experiments. In line with previous studies^79–81^, DRIP-seq peaks in wild-type (WT) and *A3B* knockout (KO) MCF10A cells were mainly intragenic and distributed between protein-coding, long non-coding RNA, and enhancer RNA genes (**Fig. 5a-b**). As anticipated by these distributions and the transcription dependence above, the vast majority of DRIP-seq positive regions in WT and KO MCF10A cells occurred in expressed genes (**Fig. S3a**).

**Fig. 5.**
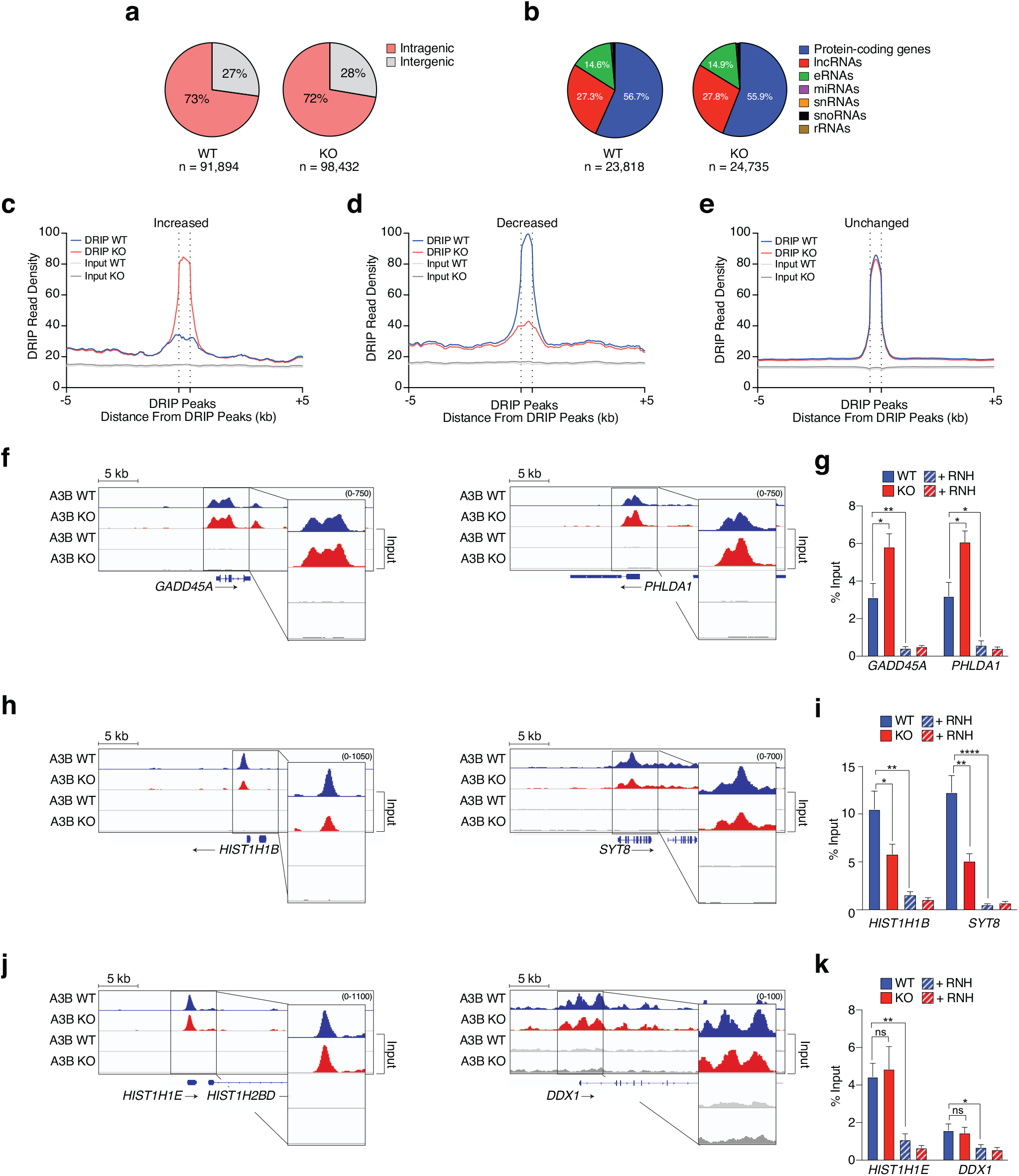
A3B affects a large proportion of R-loops genome-wide. **a-b**, Pie graphs representing R-loop distributions in wild-type (WT) and *A3B*-null (KO) MCF10A cells. **c-e**, Meta-analysis of read density (FPKM) for DRIP-seq results from wild-type (WT, blue) and *A3B*-null MCF10A (KO, red) partitioned into 3 groups (increased, decreased, and unchanged) as described in the text. Input read densities are indicated by overlapping gray lines. **f-g**, **h-i**, **j-k**, DRIP-seq profiles for representative genes in each of the groups defined in panel c-e. Independent quantification by DRIP-qPCR -/+ RNase H (RNH) treatment (striped bars) is shown in the histogram to the right (n ≥ 4; means ± SEM expressed as percentage of input; *, *P* < 0.05; **, *P* < 0.01; ****, *P* < 0.0001 by two-tailed unpaired t-test; ns, not significant).

A global comparison of DRIP-seq peaks between KO and WT MCF10A revealed changes in the overall R-loop landscape with 8,296 peaks ‘increased’, 13,761 peaks ‘decreased’, and 154,036 peaks ‘unchanged’ (red vs blue traces in **Fig. 5c-e**). Representative individual gene results are shown for *GADD45A* and *PHLDA1*, *HIST1H1B* and *SYT8*, and *HIST1H1E* and *DDX1*, which show increased, decreased, and unchanged R-loop levels, respectively, in KO cells in comparison to WT cells (**Fig. 5f, h, j**). These DRIP-seq results were confirmed by gene-specific DRIP-qPCR (**Fig. 5g, i, k**). As above, DRIP-qPCR signals were reduced significantly by RNase H treatment, further confirming R-loop specificity (**Fig. 5g, i, k**, striped bars). Differential DRIP signals were not due to transcription differences between the KO and WT cell lines (**Fig. S3b**). As expected, negligible DRIP signals were found in non-expressed genes and intergenic loci (*e.g*., *TFF1* in **Fig. S3c-d**). In addition, similar DRIP results were obtained in a different cell line (*A3B*-depleted HeLa) analyzed by immunoblotting and qPCR (**Fig. S3e-g**).

### APOBEC3B accelerates the kinetics of R-loop resolution

Transcriptional activation by different signal transduction pathways is known to trigger large increases in R-loop formation^82,83^. As the datasets above were generated under normal growth conditions, an additional set of experiments was done to address whether A3B may also affect signal transduction-induced R-loops. Therefore, WT and KO MCF10A lines were treated with PMA to induce the protein kinase C (PKC) and non-canonical (nc)NF-κB signal transduction pathways and, as such, activate the transcription of many genes including endogenous *A3B*^58,84^.

First, global R-loop distributions were analyzed by DRIP-seq in WT MCF10A following 2 hrs PMA treatment in comparison to a DMSO control treatment (workflow in **Fig. S4a**). As anticipated, PMA caused changes in the overall R-loop landscape with 13,422 peaks increased, 16,432 peaks decreased, and 171,322 unchanged (**Fig. S4b-d**). Many genes such as *JUNB* and *FOS* were induced strongly by PMA treatment and showed significant increases in DRIP signals (**Fig. S4e-f**). Other genes such as *NAXE* and *ARL4D* showed decreases in DRIP signals (**Fig. S4g-h**), whereas the majority such as *GAPDH* and *GEMIN7* showed no changes in R-loop formation (**Fig. S4i-j**).

Second, a parallel set of ChIP-seq experiments was done in MCF10A with Dox-inducible A3B-eGFP (**Fig. 1c**) to assess whether any of these R-loop categories might be bound by this deaminase (workflow in **Fig. S4a**). An epitope-tagged protein was necessary because ChIP-grade antibodies have yet to be developed for endogenous A3B. Interestingly, A3B-eGFP appeared to bind preferentially to genomic DNA regions coincident with DRIP-seq peaks increased in PMA-treated cells in comparison to vehicle control treated cells (ChIP-seq results superimposed over DRIP-seq results in **Fig. S4b**). In contrast, A3B-eGFP ChIP-seq peaks were mainly unperturbed in genomic DNA regions in which R-loop levels decreased or remained unchanged upon PMA treatment (**Fig. S4c-d**). Representative results were confirmed independently by ChIP-qPCR (**Fig. S4k**). Moreover, quantification indicated that 43% of A3B ChIP peaks overlap with R-loops peaks induced by PMA (*P* = 0.0001, two-tailed binomial test; 54% overlap for ChIP peaks with >5-fold enrichment compared to uninduced Dox^-^ condition, *P* = 0.0008). These results combined to indicate that A3B binds preferentially to genomic DNA regions with evidence for R-loop accumulation.

Third, the PMA-inducibility of this system enabled an assessment of the kinetics of R-loop resolution in the presence and absence of A3B. WT and KO cells were treated with PMA or DMSO for 2 and 6 hrs and then analyzed by DRIP-seq (workflow in **Fig. 6a**). These timepoints were chosen for analysis because transcription induction and R-loop formation are rapid (2 hrs data above) and A3B protein levels are upregulated maximally by 6 hrs post-PMA treatment^58^ (**Fig. 6b-c**). For example, *JUNB* and *DUSP1* have R-loop peaks that are strongly induced by PMA at 2 hrs and these return to near-control levels by 6 hrs (**Fig. 6d**). In contrast, R-loop peaks in the same genes remained significantly elevated and/or showed delayed resolution kinetics in KO cells after 6 hrs PMA treatment (**Fig. 6d**). PMA non-responsive genes used as controls did not show R-loop induction or major differences in R-loop levels after 6 hrs PMA treatment (*e.g.*, *GAPDH* and *HSPA8* in **Fig. 6e**). Moreover, independent IF confocal microscopy studies of the kinetics of R-loop resolution following PMA treatment of WT MCF10A cells showed that nucleoplasmic R-loop levels peak at 2 hrs and decline substantially by 6 hrs, whereas no decline (even a modest increase) was observed in *A3B* KO cells (**Fig. 6f-g**). As above, all DRIP-qPCR and nucleoplasmic R-loop signals were sensitive to RNase H treatment indicating specificity (**Fig. 6d-g**). Taken together with the ChIP data above, the results of these PMA time course experiments indicated that nuclear A3B is recruited to PMA-induced R-loops and contributes to their timely resolution.

**Fig. 6.**
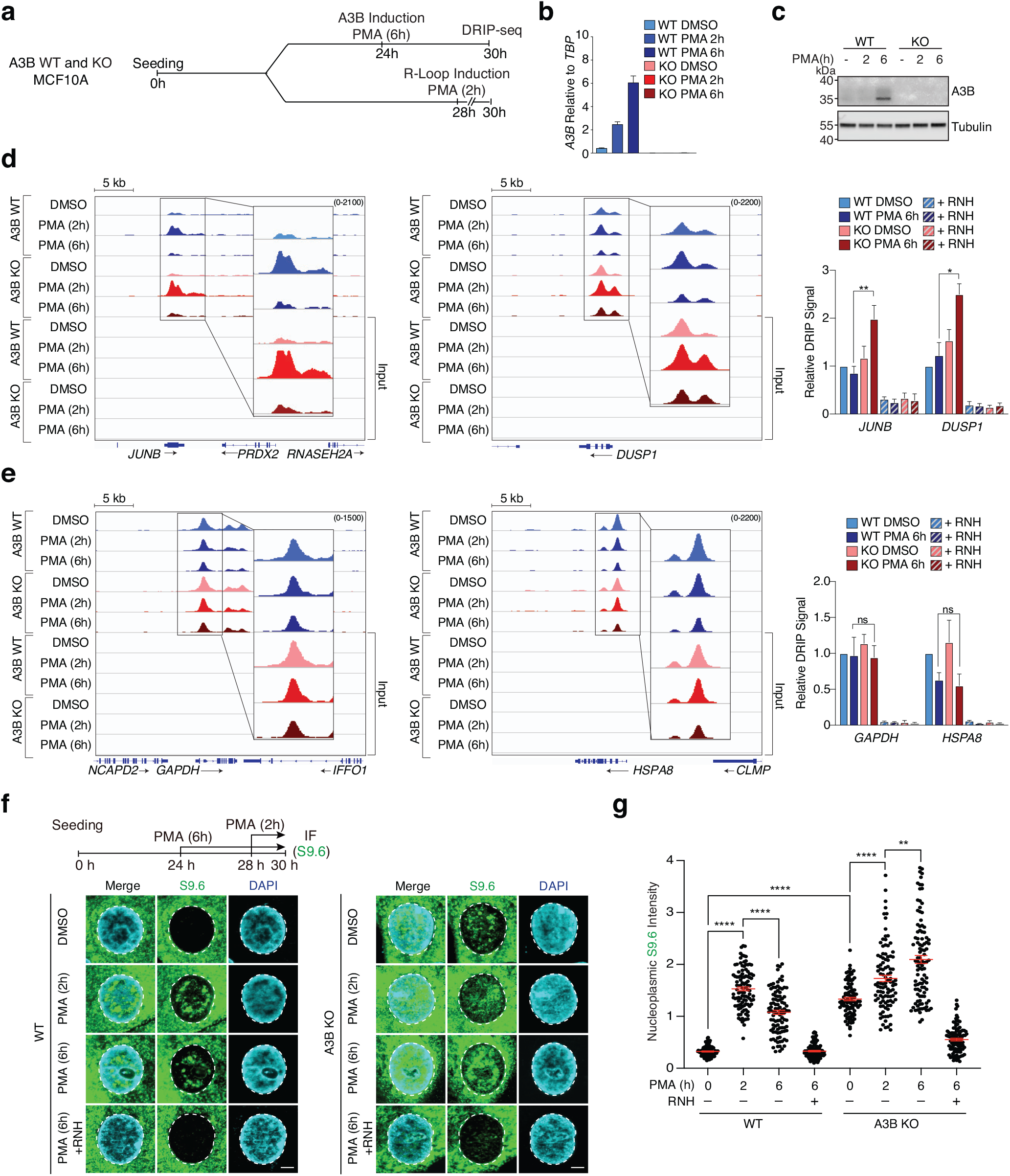
Kinetics of R-loop induction and resolution. **a**, Schematic of the DRIP-seq workflow used for panels d-e. **b**, RT-qPCR of *A3B* mRNA from MCF10A (WT) and *A3B*-null (KO) cells treated with PMA (25 ng/ml) for the indicated times. Values are expressed relative to the housekeeping gene, *TBP* (n = 3; mean ± SEM; knockout levels not detectable). **c**, Immunoblots of extracts from MCF10A (WT) and *A3B*-null (KO) cells treated with PMA (25 ng/ml) for the indicated times and probed with indicated antibodies. **d**, DRIP-seq profiles for two PMA-responsive genes, *JUNB* and *DUSP1*, in DMSO or PMA-treated MCF10A (WT; upper profiles) and *A3B*-null (KO; lower profiles). Independent quantification by DRIP-qPCR -/+ RNase H (RNH) treatment (striped bars) is shown in the histogram to the right (n ≥ 4; mean ± SEM normalized to DMSO WT; *, *P* < 0.05 and **, *P* < 0.01 by two-tailed unpaired t-test). **e**, DRIP-seq profiles for two PMA non-responsive genes, *GAPDH* and *HSPA8*, in DMSO or PMA-treated MCF10A (WT; upper profiles) and *A3B*-null (KO; lower profiles). Independent quantification by DRIP-qPCR -/+ RNase H (RNH) treatment (striped bars) is shown in the histogram to the right (n ≥ 4; mean ± SEM normalized to DMSO WT; ns, not significant). *HSPA8* levels decline in WT and KO cells after PMA treatment in a manner that is independent of A3B. **f-g**, IF images and quantification of MCF10A (WT) and *A3B*-null (KO) cells treated with PMA for the indicated times and stained with S9.6 (green) and DAPI (blue). Diminution of nucleoplasmic S9.6 signal by exogenous RNase H (RNH) confirmed signal specificity (representative images; 5 μm scale bar; n > 100 nuclei per condition; red bars represent mean ± SEM; **, *P* < 0.01 ****, *P* < 0.0001 by Mann-Whitney test).

### APOBEC3B biochemical activities required for R-loop resolution

To investigate the biochemical activities of A3B in R-loop resolution, wild-type A3B was purified from 293T cells (**Fig. S5a**) and used for nucleic acid binding and DNA deamination experiments (**Fig. 7** and **Fig. S5b-c**). The various nucleic acids used in these biochemical experiments are depicted in **Fig. 7a**. EMSAs indicated that A3B is capable of binding to R-loop structures (novel), to ssDNA and ssRNA (expected^67,68,85^), and to lesser extents to dsDNA, dsRNA, or RNA/DNA hybrid (also expected^67,68,85^; **Fig. S5b-c**). These native EMSAs were hard to quantify due to accumulation of large protein/nucleic acid complexes in the wells. We therefore reduced the complexity of binding studies by quantifying the release of fluorescently-labeled ssRNA and ssDNA from A3B by incubating with a broad concentration range of otherwise identical unlabeled nucleic acid competitors. These experiments demonstrated that A3B binds strongly and nearly indistinguishably to both ssRNA and ssDNA (**Fig. 7b**).

**Fig. 7.**
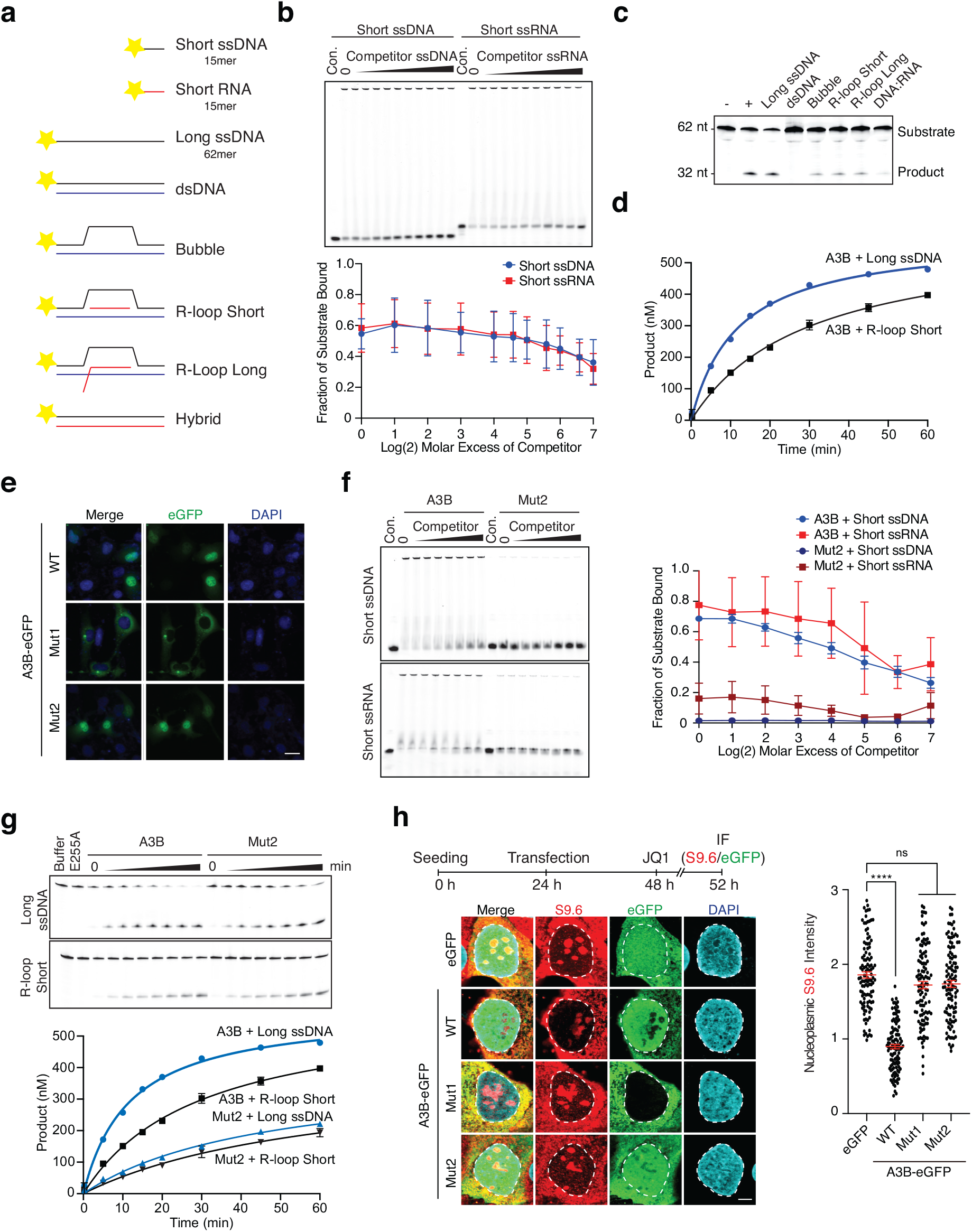
A3B biochemical activities required for R-loop resolution. **a**, Schematics of the nucleic acids used in biochemical experiments (5’ fluorescent label indicated by yellow star). The 15mer short ssDNA and short RNA were used in EMSAs in panels b and f, and the 62mer long ssDNA was used alone or as annealed to the indicated complementary nucleic acids (black, DNA; red, RNA) in other experiments. **b**, Native EMSAs of A3B binding to fluorescently labeled short 15mer ssDNA or RNA in the presence of increasing concentrations of otherwise identical unlabeled competitor. The corresponding quantification shows the average fraction bound to substrate +/- SD from 3 independent experiments. **c**, Substrates in panel a tested qualitatively for deamination by A3B. Negative (-) and positive (+) controls are the long ssDNA alone and deaminated by recombinant A3A. **d**, A quantitative time course of A3B-catalyzed deamination of the long ssDNA vs an R-loop (short) substrate (mean +/- SD of n = 3 independent experiments are shown with most error bars smaller than the symbols). **e**, Subcellular localization of A3B-eGFP (WT), Mut1, and Mut2 in U2OS cells (scale bar = 10 µM). **f**, EMSAs comparing A3B WT and Mut2 binding to short 15mer ssDNA and RNA in the presence of increasing concentrations of otherwise identical unlabeled competitor ssDNA or RNA. The corresponding quantification shows the average fraction bound to substrate +/- SD from 3 independent experiments. **g**, Quantitative comparison of A3B WT and Mut2 deamination of the long ssDNA versus an R-loop (short) substrate. Representative gels are shown for the time-dependent accumulation of product, along with quantitation of 3 independent experiments (mean +/- SD with most error bars smaller than the symbols; for comparison the WT data are the same as those in panel d). **h**, IF images and quantification of U2OS cells expressing the indicated eGFP construct (green) and stained with S9.6 (red) and DAPI (blue; 5 μm scale bar; n > 60 nuclei per condition; red bars represent mean ± SEM; ****, *P* < 0.0001 by one-way ANOVA; ns, not significant).

RNA is an inferred inhibitor of the ssDNA cytosine deaminase activity of A3B based on experiments in which exogenous RNase A treatment is required to detect ssDNA deaminase activity in cancer cell extracts (*e.g*., refs.^22,58^). We therefore wondered whether the RNA in R-loop structures might inhibit the ssDNA deaminase activity of A3B on the unpaired ssDNA strand. Qualitative single timepoint reactions indicated clear activity on free ssDNA cytosines and potentially reduced activities on ssDNA cytosines in a bubble, short R-loop, and long R-loop structures (**Fig. 7c**). A quantitative time course comparing A3B activity on free ssDNA versus ssDNA in the context of an R-loop structure indicated that the latter substrate is only about 2-fold less preferred (**Fig. 7d**). These data showed that R-loops can be substrates for A3B-catalyzed ssDNA deamination. The 2-fold diminution in activity may be due to ssDNA inaccessibility caused by the relatively short nature of the synthetic R-loop (21 nucleotides) and/or to competition with unpaired ssDNA or ssRNA.

To gain additional mechanistic insights into A3B function in R-loop biology, we analyzed the nucleoplasmic R-loop phenotypes of an A3B mutant defective in nuclear localization (Mut1)^86^ and a mutant incapable of binding strongly to nucleic acid (Mut2)^85^. Both of these activities are governed by the N-terminal domain (NTD) of A3B, and are independent of the C-terminal domain (CTD) that binds much more weakly to ssDNA and catalyzes deamination^67,85,86^. We confirmed the nuclear localization defect of the Mut1 and, interestingly, showed that Mut2 still retains this hallmark activity (**Fig. 7e**). Mut2 was also purified and, in contrast to the wild-type enzyme, confirmed defective in binding to ssRNA and ssDNA using EMSAs (**Fig. S5a** and **Fig. 7f**). However, Mut2 still retained high levels of ssDNA deaminase activity and was similarly active on free ssDNA and short R-loop ssDNA (**Fig. 7g**). This result is consistent with the possibility that unannealed nucleic acid may interfere with the deaminase activity of the WT enzyme (above) but not with Mut2, which has reduced nucleic acid binding activity. Most importantly, in contrast to WT A3B, neither mutant was capable of decreasing nucleoplasmic R-loop levels following JQ1 treatment (**Fig. 7h**). The separation-of-function Mut2 protein also had a diminished capacity to co-IP interactors (**Fig. S1f**). These results indicated that both nuclear localization and nucleic acid binding activities are required for A3B to alter nucleoplasmic R-loop levels.

### Genome-wide associations between APOBEC3 signature mutations, gene overexpression, and splicing defects

The involvement of A3B in transcription-dependent R-loop biology led us next to evaluate whether transcribed genes may be preferred sites of mutagenesis. Specifically, we propose a working model in which exposed ssDNA cytosines in R-loop regions are deaminated by A3B and resolved into mutagenic or non-mutagenic outcomes (**Fig. 8a** and **Discussion**). Non-mutagenic outcomes are likely to comprise most resolution events, but these leave no genomic scars and, thus, are invisible to bioinformatics analyses. However, mutagenic R-loop resolution outcomes are predicted to reflect the intrinsic structural preference of A3B for deamination of cytosines in TC motifs^7,87,88^ and more broadly in TCW and RTCW extended motifs as defined experimentally and computationally^23,26^. For comparison, A3A elicits a preference for YTCW^23,26,89^.

**Fig. 8.**
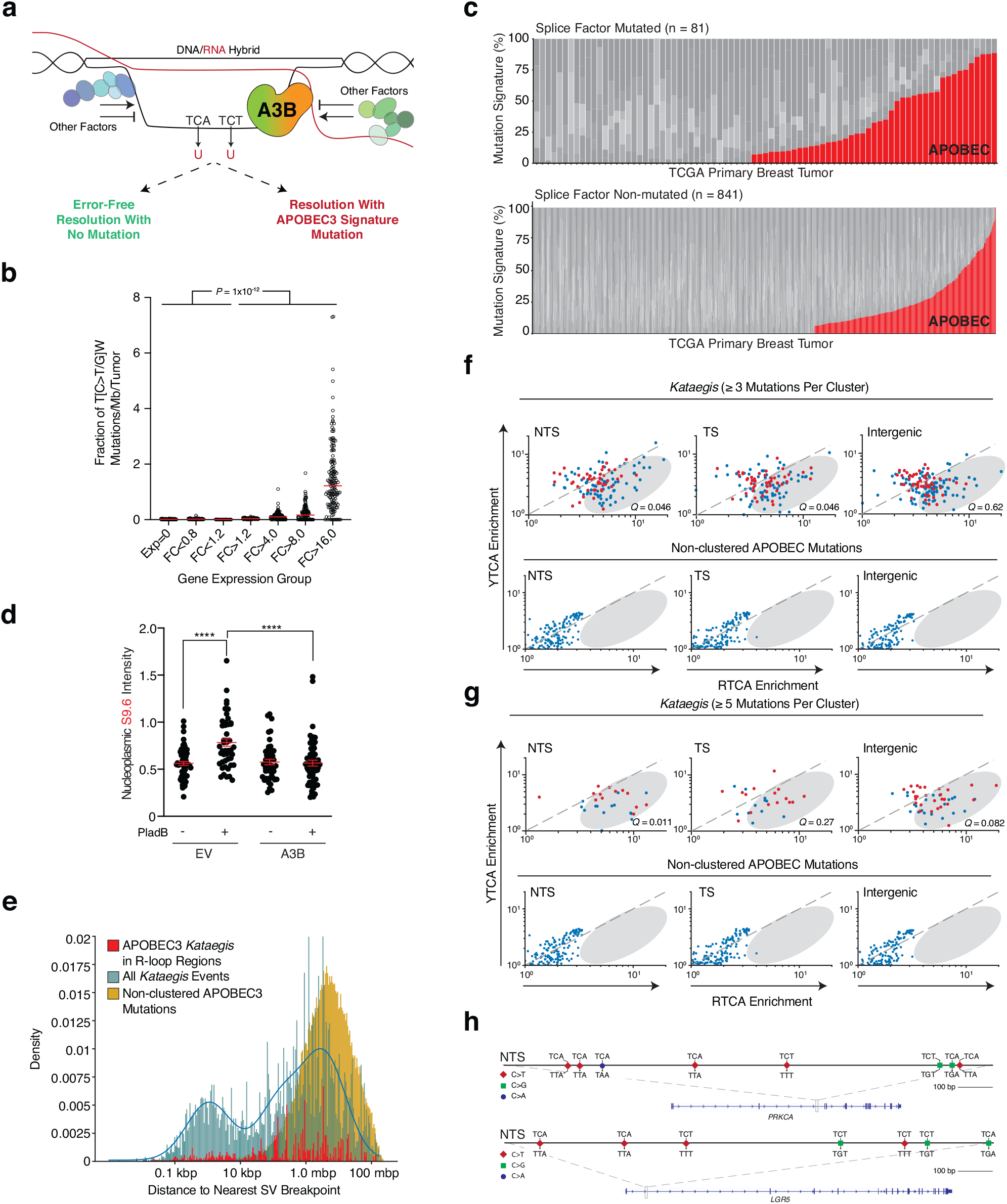
R-Loop mutagenesis and *kataegis* by APOBEC3B. **a**, Cartoon of a working model for A3B-mediated R-loop resolution with and without associated mutations. Other R-loop regulatory factors are depicted in shades of green and blue. Transcription, splicing, and other RNA- and R-loop-associated complexes are not shown for clarity. **b**, A dot plot showing the fraction of APOBEC3-attributed mutations (per Mbp per tumor) in the indicated overexpressed gene groups (FC, fold-change in breast tumors relative to the average observed in normal breast tissues). Pairwise comparisons are significant for all combinations of the lowest 4 versus the highest 3 expression groups (*P* < 1 x 10^-12^ by Welsh’s t-test). **c**, Stacked bar graphs showing the proportion of each COSMIC mutation signature in TCGA breast tumors with mutations in splice factor genes or not (n = 81 splice factor mutated tumors; n = 841 for non-splice factor mutated tumors; *P* < 0.017 by Fisher’s exact test). The APOBEC3 signature percentage (red) is comprised of COSMIC signatures 2 and 13, and other signatures are shown in different shades of gray. **d**, Quantification of nucleoplasmic R-loop levels in U2OS cells expressing an empty vector (EV) control or A3B following a 2 hr treatment with DMSO or the splicing inhibitor Plad B (n > 50 nuclei per condition; red bars represent mean ± SEM; ****, *P* < 0.0001 by two-tailed unpaired t-test). **e**, Distribution of the distances to the nearest SV of all non-clustered APOBEC3 mutations (gold), all *kataegic* mutation events (teal), and R-loop associated APOBEC3 *kataegic* mutations (red). **f-g**, Dot plot representations of short (≥3) and long (≥5) APOBEC3 *kataegic* tracts in PCAWG breast tumor WGS. The x-and y-axis reports A3B-associated 5’-RTCA and A3A-associated 5’-YTCA enrichments, respectively. Dashed lines indicate where non-skewed distributions should cluster, red dots represent data from specimens with at least one R-loop-associated *kataegic* event, and the gray ovals highlight the area in which 5’-RTCA enrichments occur (Mann-Whitney U-test *Q*-values indicated in each plot). **h**, Representative NTS *kataegic* events in the indicated genes. Wild-type trinucleotides and mutational outcomes are indicated.

One prediction of this model is that a positive association may exist between transcription levels and APOBEC3-attributed mutations. Because previous studies established a correlation between gene expression and R-loop formation^79^, we predicted that higher rates of transcription should, at least at some loci, lead to higher rates of R-loop formation and increased exposure to A3B. This idea was addressed using whole-exome sequenced (WES) breast cancers and corresponding RNA-seq data from TCGA project as well as whole-genome sequenced (WGS) breast cancers from the ICGC consortium and normal breast tissue gene expression data from the GTEx project (Methods). An initial association between gene expression levels and APOBEC3-attributed mutations was intriguing but became insignificant after accounting for gene size (**Fig. S6a-b**). However, a strong positive association emerged between the magnitude of gene overexpression in breast cancer compared to normal breast tissue and the proportion of mutations attributable to APOBEC3 deamination (all TCW mutations in **Fig. 8b**, *P* < 1.0 x 10^-12^ by student’s t-test; RTCW/YTCW breakdown in **Fig. S6c**). These analyses indicated that the higher the degree of gene overexpression in breast cancer the higher the proportion of mutations attributable to APOBEC3, with the highest overexpressed gene group showing an average of over 50-fold more APOBEC3 signature mutations than any of the three lowest expressed gene groups.

Second, because splicing defects lead to increases in R-loop formation^90–92^, our working model predicts that splice factor mutant tumors may manifest elevated levels of APOBEC3 signature mutations. This idea was investigated by splitting the TCGA breast cancer WES data set into tumors with and without mutations in splice factor genes and evaluating associations with the proportion of mutations attributable to APOBEC3 activity. Remarkably, 53% of the breast tumors with mutant splice factor genes (43/81) had significant levels of APOBEC3 signature mutations (**Fig. 8c**). In contrast, only 35% of breast tumors without mutations in the same splice factor gene set (326/841) showed a detectable APOBEC3 mutation signature (**Fig. 8c**; *P* < 0.017 by Fisher’s exact test). APOBEC3-attributed mutations in both groups were partitioned into RTCW/YTCW tetranucleotide motifs and, interestingly, the A3B-associated RTCW motif was only absent from one of the splice factor mutant tumors (1/43) in comparison to a significant proportion of the non-splice factor mutant group (52/326) (**Fig. S6d-e**; *P* = 0.028 by Fisher’s exact test). Splice factor mutant tumors also had a higher mean percentage of APOBEC3-attributed mutations (39% vs 31%, respectively; *P* = 0.042 by unpaired two-sample Welsh’s t-test) as well as higher total numbers of mutations on average than non-splice factor mutated samples (*P* = 0.0018 by Welch’s two sample t-test). Even the top quartile of tumors with the strongest APOBEC3 signature had higher total numbers of mutations in the splice factor mutant group (*P* = 0.0095 by Welch’s two sample t-test). These relationships between splice factor defects, higher mutation loads, and APOBEC3 mutation signature are not likely due to chance because no other similarly sized gene set selected randomly from housekeeping genes (100,000 random gene set selections) is similarly mutated in TCGA breast cancer data sets (*i.e*., the observed splice factor defects are not due to higher rates of mutation; rather, the observed splice factor defects are likely to contribute to the higher rates of mutation). We also noted that APOBEC3-attributed mutations accumulate preferentially on the non-transcribed strand (NTS) over the transcribed strand (TS) in both splice factor mutant and non-mutant tumor groups with a statistical difference that may relate to the underlying mechanism (*P* = 0.0284 and *P* = 4.3 x 10^-12^, respectively, by student’s t-test). In further support of a connection between aberrant splicing, R-loop formation, and APOBEC3 mutagenesis, A3B overexpression suppresses the increase in R-loop formation caused by treating U2OS cells with the splicing inhibitor pladienolide B (Plad B; **Fig. 8d**).

Third, because APOBEC3 signature *kataegic* events are undoubtedly due to strand-coordinated deamination by at least one APOBEC3 enzyme^20,25^, we asked what proportion of these events occur in genes and, moreover, occur on the NTS versus the TS to potentially reveal which part of the genome may be more susceptible. This analysis focused again on breast cancer due to the large size and overall high quality of the WGS data sets. A global mapping of all *kataegis* events in primary breast adenocarcinomas from the Pan-Cancer Analysis of Whole Genomes (PCAWG) revealed a bimodal distribution with one peak located within 1 kbp of a structural variation breakpoint (SV) and another similarly sized peak much further away from the nearest SV (∼1 Mbp; n = 198 WGS data sets; blue bars in **Fig. 8e**). As expected^93^, the SV-proximal subset of *kataegis* events is likely due to deamination of resected ssDNA ends during recombination repair. Also expected, non-clustered (*i.e*., dispersed) APOBEC3-attributed mutations occur on average of >1 Mbp apart (yellow bars in **Fig. 8e**). In contrast, the majority of APOBEC3-attributed *kataegic* events (>75%) map >10 kbp away from SVs and are unlikely to be due to a DNA double-stranded break and recombination-associated mechanism (red bars in **Fig. 8e**). Moreover, a significant proportion of these distal APOBEC3 signature *kataegic* events occur within R-loop regions defined above in DRIP-seq experiments.

To investigate the source of APOBEC3 signature *kataegis*, we plotted overall enrichments for A3B-associated RTCA and A3A-associated YTCA tetranucleotide motifs (R=A or G; Y=C or T)^26^ and shaded data points red to distinguish breast tumors with at least one R-loop associated *kataegic* event. This analysis indicated, first, that tumors with APOBEC3 signature *kataegis* are common and, second, that APOBEC3 *kataegic* mutations overlapping R-loop regions, in contrast to dispersed APOBEC3 mutations, are skewed toward RTCA motifs (**Fig. 8f-g**; *Q*-values in each dot plot determined using Mann-Whitney U-tests). Third, the overall RTCW skew of *kataegic* (≥3 mutations per cluster) versus dispersed APOBEC3 mutations was greatest for mutations occurring on the NTS but still evident on the TS and intergenic regions (**Fig. 8f**; *P* = 0.0006, *P* = 0.0086, and *P* = 0.096 respectively, by Fisher’s exact tests after FDR correction). For greater stringency, this latter analysis was repeated for longer APOBEC3 *kataegic* tracts (≥5 mutations per cluster) and a statistically significant enrichment was only evident for RTCA events on the NTS of genes [**Fig. 8g**; *P* = 0.0001 (NTS), *P* = 0.15 (TS), and *P* = 0.072 (intergenic)]. Representative NTS *kataegic* events are shown for *PRKCA* and *LGR5* (**Fig. 8h**). Taken together, these different bioinformatic analyses support a model in which at least a subset of R-loop structures is susceptible to C-to-U deamination events most likely catalyzed by A3B.

## Discussion

Our studies are the first to report the cellular interactome of A3B and reveal an unanticipated role for this antiviral enzyme in R-loop biology. We delineate a functional relationship between A3B and R-loops with higher R-loop levels occurring upon A3B knockout/down and lower R-loop levels upon A3B overexpression. Genome-wide DRIP-seq experiments in physiological conditions and upon activation of a signal transduction pathway indicated that thousands of R-loops in cells are affected by A3B (*i.e.*, increased or decreased in *A3B*-null cells). This number represents over 10% of R-loops genome-wide, which is comparable to the proportions of R-loops affected by other R-loop regulatory factors, such as TOP1, DDX5, XRN2, and PRMT5^80,94^. These findings are also in line with knowledge that multiple proteins contribute to R-loop regulation, including RNase H1, TOP1, SETX, AQR, UAP56/DDX39B, FANCD2, and BRCA1/2^38,45,80,94–103^. Determining the precise combination of molecular mechanisms responsible for the regulation of a given R-loop remains a challenge for future studies.

In addition to discovering an unanticipated role for A3B in R-loop homeostasis, our studies also shed light on the underlying molecular mechanism of R-loop resolution. First, complementation experiments showed that the mechanism is deaminase-dependent with expression of a single amino acid catalytic mutant (E255A) failing to decrease nucleoplasmic R-loop levels. Second, the nuclear localization activity of A3B is essential. This requirement may seem obvious but is important to help rule-out potential indirect effects such as A3B binding to cytoplasmic factors and affecting their import into the nuclear compartment and participation in R-loop formation. Third, A3B is capable of binding to R-loop structures, and the strong nucleic acid binding activity of A3B is required for suppressing nucleoplasmic R-loop levels. Our biochemical competition experiments indicated that ssRNA and ssDNA binding activities are comparable in strength. Together with the fact that A3B’s strong nucleic acid binding activity resides within the N-terminal half of the protein and the weaker ssDNA binding activity required for catalysis is governed by the C-terminal half of the enzyme, we favor a working model in which direct binding of A3B to nascent ssRNA adjacent to R-loops and/or to ssDNA exposed in R-loop structures is critical for R-loop regulation (**Fig. 8a**). The next steps in such a mechanism may be deamination of exposed ssDNA cytosines in R-loop structures, uracil excision by UNG2, ssDNA breakage by APEX, and recruitment of additional downstream repair factors for resolution, by analogy to the resolution of AID-catalyzed R-loop associated deamination events in immunoglobulin gene switch regions^41,42^ and APOBEC-catalyzed editing of R-loop DNA cytosines exposed in Cas9-mediated base editing reactions^104^. Additional tests of this model should consider complications from synthetic lethal interactions such as between A3B overexpression and UNG2 inhibition^52^. Moreover, the relationship between R-loop homeostasis and DNA repair is likely to be complex, and it is easy to imagine how perturbing R-loop levels in either direction (up or down) could lead to elevated DNA damage responses (*e.g*., γ-H2AX accumulation), as observed here with A3B-deficient cells and elsewhere with other factors^80,95,96,99,105,106^. Notably, forced overexpression of A3B can also cause elevated DNA damage responses including γ-H2AX accumulation^22,52,107^.

This proposed mechanism is likely to lead to predominately error-free resolution, which cannot be detected by bioinformatic genomic analyses. However, a subset of the uracil lesions and/or abasic sites may lead to APOBEC3 signature mutations and, if clustered, to APOBEC3 *kataegic* events. The possibility of such mutagenic outcomes is supported by a dose-responsive association between genes overexpressed in breast cancers compared to normal breast tissue and the proportions of somatic mutations attributed to APOBEC3 deamination (**Fig. 8b**). Moreover, a preferential accumulation of APOBEC3-attributed mutations is also observed in splice factor mutant breast tumors (**Fig. 8c**). As transcription and splicing are interconnected processes^108^ and splice factor defects are known to increase R-loop levels^90–92,109^, it is easy to imagine how perturbations in these processes may contribute to increased R-loop formation and exposures of single-stranded cytosines to APOBEC3 deaminase activity. This possibility is further supported by recent bioinformatic studies indicating higher APOBEC3 mutation densities on the NTS of actively expressed genes (including the overexpressed genes analyzed here) in multiple cancer types^110^. We cannot exclude the possibility that other APOBEC3 enzymes, most notably A3A, may also be able to contribute to R-loop mutation. However, this alternative is disfavored because A3A overexpression does not affect R-loop levels and, importantly, the majority of APOBEC3 *kataegic* events are observed far away from sites of structural variation and are enriched for mutations in A3B-associated 5’-RTCW motifs (**Fig. 8e-h**).

In addition, because a proportion of APOBEC3 mutagenesis is likely due to deamination of lagging-strand templates during DNA replication^29–31,33,34,94^, it is important to emphasize that APOBEC3-catalyzed C-to-U lesions in transcription-associated R-loop structures are likely to attract different subsets of DNA repair enzymes. A uracil in the DNA replication template strand will instruct the proper insertion of an adenine in the nascent strand, which leaves a U:A base pair for UNG2 recognition and faithful uracil base excision repair with the net result being a C-to-T transition. A uracil in an R-loop, following R-loop resolution, will lead to a U/G mispair, which may similarly template the insertion of an adenine through replication but can also attract mismatch repair enzymes, which could promote the creation of a longer ssDNA tract and additional APOBEC3 mutagenesis. This possibility is supported by recent results showing a mismatch repair-dependent synthetic lethal interaction between A3B activity and uracil excision repair disruption^52^ and a role for mismatch repair in the creation of *omikli* (shorter-than-*kataegic* tracts of APOBEC3-attributed mutations)^111^. It is also supported by precedents with AID-dependent antibody gene diversification in B lymphocytes where both uracil base excision repair and mismatch repair have integral roles^41,42^. Of course, uracil lesions in both DNA replication template strands and R-loops can lead to alternative outcomes including uracil excision, abasic site bypass or cleavage, single- and double-stranded DNA breaks, and larger-scale genetic aberrations including translocations. A3B-mediated mutagenesis may be particularly relevant in cancer cells that display both increased A3B expression and R-loop dysregulation resulting from mutations in splice factors or other R-loop regulatory factors (*e.g.*, BRCA1/2 and Fanconi anemia proteins^99,100,102,112,113^) and/or activation of signal transduction pathway-induced transcriptional programs^82,83^ such as during conditions of oncogenic signaling, infection, or inflammation^58,114,115^. Therefore, in addition to archetypic base substitution mutations, it is tempting to speculate that at least some R-loop-associated DNA damage and chromosome aberrations (reviewed by refs.^39,46,48,116^) may be instigated by the A3B-dependent mechanism described here.

In addition to the resolution mechanism discussed above, a potentially overlapping alternative is A3B-dependent recruitment of other proteins known to promote R-loop resolution. Such interactions could be direct or bridged, for instance, by RNA or ssDNA. In support of this possibility, the A3B separation-of-function mutant Mut2, which is deficient in nucleic acid binding but proficient in nuclear import and DNA deamination, is less capable of interacting with several R-loop associated factors (Fig. S1f). Moreover, although our studies here focused on AP-MS interactions specific to A3B (absent from all controls), a number of A3B-enriched factors (present at lower levels in at least one control reaction), such as the helicase DHX9, might be worthy of future investigation. This DNA/RNA helicase was reported recently as a suppressor of the antiviral activity of A3B^117^. It is harder to envisage a mechanism to explain the subset of genes that elicit decreased R-loop levels in the absence of A3B. In these instances, A3B may be blocking a known resolution activity, such as a helicase that loads via RNA or ssDNA, from gaining access to R-loops. Further studies on A3B regulation of R-loop homeostasis will undoubtedly be exciting as answers will shed additional light on R-loop biology, provide insights into the normal physiological functions of A3B in innate antiviral immune responses, and may help define additional targetable nodes in A3B-overexpressing tumor types such as breast cancer.

## Acknowledgements

We thank N.J. Proudfoot for critically reading the manuscript, J. Becker and J. Duda for corroborative localization data with A3B mutants, the University of Minnesota Imaging Center for access to instrumentation, and the Oxford Genomics Centre at the Wellcome Centre for Human Genetics (funded by Wellcome Trust grant 203141/Z/16/Z) for the generation and initial processing of the sequencing data. This work was supported by NCI P01 CA234228 (to RSH), NIAID R37 AI064046 (to RSH), and by the University of Minnesota Masonic Cancer Center, Academic Health Center, and College of Biological Sciences. NG lab is supported by the Royal Society University Research fellowship (BVD07340), Royal Society Enhancement Award (RGF\EA\180023) and EPA Research Fund (Sir William Dunn School of Pathology, University of Oxford) to NG and CRUK development fund (CRUK DF-0119) to AC and NG. MT and SM are supported by the Wellcome Trust Investigator Award to SM (WT210641/Z/18/Z). KMM lab was supported by NCI RO1 CA198279 and NCI RO1 CA201268. LBA lab was supported by US NIH R01 ES030993 and R01 ES032547. Salary support for JLM was provided initially by an NSF Graduate Research Fellowship (Grant Number 00039202) and subsequently by HHMI. Salary support for MCJ was provided in part by T32 CA009138 and subsequently NCI F31 CA243306. Salary support for BS was provided by HHMI with some supplies provided by the Ovarian Cancer Research Alliance (Ann and Sol Schreiber Mentored Investigator Award 812337). Salary support for DJS was provided by NIAID K99 AI147811. RSH is the Margaret Harvey Schering Land Grant Chair for Cancer Research, a Distinguished University McKnight Professor, and an Investigator of the Howard Hughes Medical Institute.

## Author contributions

R.S.H., J.L.M., A.C. and N.G. conceived and designed these studies. J.L.M. and A.C. performed experiments unless otherwise noted. E.K.L. generated U2OS knockdown and complement cell lines along with assisted in tissue culture and genomic DNA isolations for dot blot experiments. S.L., J.K. and K.M.M. performed and quantified IF data. M.T. and S.M. conducted DRIP-seq and ChIP-seq data analysis. C.B. performed DRIP-qPCR validations and HeLa R-loop IP. B.S. assisted with cell culture experiments and R-loop quantification. M.R.B. assisted with cell culture studies. M.C.J., N.A.T., D.J.S., E.N.B. and L.B.A. performed bioinformatic analyses. M.A.C. contributed biochemical experiments. R.S.H. and J.L.M. drafted the manuscript with input from all other authors.

