## Supplemental methods, Table 2, and Figures S1-S6 for "R-loop homeostasis and cancer mutagenesis promoted by the DNA cytosine deaminase APOBEC3B"

**SI contents:**

- Methods
- Tables S1-2
- Figs. S1-6

### Methods

**Cell lines and culturing.** U2OS cells were obtained from ATCC (ATCC HTB-96) and were maintained in McCoy's 5A Medium (ThermoFisher, 16600082) supplemented with 10% FBS (Gibco, Gaithersburg, MD) and 0.5% pen/strep (50 units). U2OS shCtrl and shA3B cell lines were made using previously described shCtrl and shA3B lentiviral constructs, viral production and transduction methods and puromycin selection 1 µg/mL[1]. U2OS pcDNA3.1-A3-3xHA stable lines were made via linear (*Nru*I digested) transfection and selection using 800 µg/mL G418. HEK 293T cells were obtained from ATCC (#CRL-3216) and were maintained in RPMI (Hyclone, South Logan, UT) supplemented with 10% FBS (Gibco, Gaithersburg, MD) and 0.5% pen/strep (50 units). MCF10A cells were obtained from ATCC (ATCC CRL-10317) and were maintained in DMEM/F12 (Invitrogen, 11330-032) supplemented with 5% horse serum (Invitrogen, 16050-122), 20 ng/mL EGF (Peprotech), 0.5 mg/mL hydrocortisone (Sigma Aldrich, H-0888), 100 ng/mL cholera toxin (Sigma Aldrich, C-8052), 10 $\mu$g/mL insulin (Sigma Aldrich, I-1882; Sigma Aldrich, I-9278) and 0.5% pen/strep (Invitrogen, 15070-063). MCF10A-TREx-A3B-eGFP were maintained in the same MCF10A media described above with the addition of 100 µg/mL Normocin. S9.6 Hybridoma cells were obtained from ATCC (ATCC HB-8730) and were maintained in DMEM (Hyclone, South Logan, UT) supplemented with 10% FBS (Gibco, Gaithersburg, MD) and 0.5% pen/strep (50 units). HeLa cells were obtained from ATCC and were maintained in DMEM (Sigma-Aldrich) supplemented with 10% FBS (Sigma-Aldrich) and 0.5% pen/strep (Invitrogen, 15070-063). MCF10A *A3B* KO cell line was engineered by transduction with pLentiCRISPR, expressing the gRNA sequence GCTCCATTCAACCCCCCTGCT targeting both the *A3A* and *A3B* genes. Cells were selected with puromycin and seeded for single cell cloning. Deletion mutant lines were identified by PCR using primers amplifying unique sequences within the *A3B* gene and/or the *A3A/B* junction (primers in ref.[2]) and confirmed by qPCR and immunoblots. HeLa RNAi was performed in 6-well plates 24 hrs after seeding with 22 nM siRNA and Lipofectamine 2000 and after 6hrs the medium was changed. A second transfection was performed 48 hrs after seeding using the same experimental setting and then, cells were re-seeded 24 hrs before the experiment. siRNAs were purchased from GE Healthcare targeting Luciferase (D-001400-01) or APOBEC3B (CCUGAUGGAUCCAGACACAdTdT).

**Plasmids and cloning.** C-terminal eGFP epitope-tagged plasmids used in this study were described previously[1,3-5]. Catalytic mutant A3B-E255A and shRNA-resistant derivatives were made using standard site-directed-mutagenesis. C-terminal 3x-HA epitope-tagged plasmids used in this study were described previously[6] and shRNA-resistant derivatives were made using standard site-directed-mutagenesis. C-terminal 2xStrep and 3xFlag-tagged eGFP and A3B constructs used for proteomics were described previously[7]. cDNA for some interactors constructs were ordered from Origene (RC216648, RC204785 and RC214037) while the rest were cloned from 293T cDNA. 4/TO-C-terminal 3xFlag-tagged interactor constructs used for immunoprecipitation were generated using standard cloning techniques. A3B Mut1(E22Y/E24R/Y28S/G29R/S31N/Y32T)[8] and Mut2 (Y13D/Y28S/Y83D/W127S/Y162D/Y191H)[9] were subcloned into 5/TO A3B-GFP as a *Hind*III and *Kpn*I fragment from a reported construct or gBlock (IDT), respectively. All oligonucleotide sequences used to generate new constructs are listed in **Table S2**.

**Affinity purification and mass spectrometry.** HEK 293T cells were transfected with pcDNA4/TO-A3B-2xStrep-3xFlag or eGFP-2xStrep-3xFlag using Transit LT1 (Mirus). Cells were harvested in 1x PBS 48 hrs post-transfection. Cells were washed two times in 1x PBS followed by lysis (50 mM Tris-HCl pH 8.0, 1% Tergitol NP-40, 150 mM NaCl, 0.5% sodium deoxycholate, 0.1% SDS, 1 mM DTT, 1x Protease Inhibitor [Roche], RNase A and DNase). Lysates were subjected to sonication prior to clearing by centrifugation. Cleared lysates were then added to Strep-Tactin Superflow resin (IBA) followed by end-over-end rotation for 2 hrs at 4°C. Following IP, the anti-Strep resin was washed three times in high salt wash buffer (20 mM Tris-HCl pH 8.0, 1.5 mM $MgCl_2$, 1 M NaCl, 0.2% Tergitol NP-40, 0.5 mM DTT and 5% glycerol) followed by three washes in low salt wash buffer (same as high salt but with 150 mM NaCl). To remove detergents for proteomics submission, samples were subjected to three washes of no-detergent wash buffer (20 mM Tris-HCl pH 8.0, 1.5 mM $MgCl_2$, 150 mM NaCl, 0.5 mM DTT and 5% glycerol). Protein was eluted from the resin in elution buffer (100 mM Tris-HCl pH 8.0, 150 mM NaCl and 2.5 mM desthiobiotin). Samples were validated using immunoblotting, DNA deaminase activity assays (discussed below), and Coomassie staining. In-solution samples were analyzed by LC-MS/MS at the Harvard Proteomic Core (data are summarized in **Table S1**). CRAPome repository was used to remove likely non-specific interactions prior to S9.6 IP overlap analysis[10].

For A3B-mycHis purification, 293T cells grown in RPMI were transfected in 15 cm plates with 20 μg of plasmid using a 3:1 ratio of polyethyleneimine (Polysciences

Inc. PEI 40k #24765) to DNA. 24 hours post-transfection, the cells were harvested by trypsinization, washed in PBS-EDTA, and collected by centrifugation. Cell pellets were frozen at -80 °C. For purification, cells were lysed in 25 mM HEPES pH 7.4, 300 mM sodium chloride, 20 mM imidazole, 10 mM magnesium chloride, 0.5 mM TCEP, 0.1% Triton X-100, 20% glycerol, and Roche Complete Protease inhibitors. Lysis was performed by two minutes of sonication at a 40% duty cycle. Following sonication, RNase A was added to 100 µg/mL and Benzonase to 5 units/mL followed by incubation at 37 °C for an hour. Cell debris was pelleted by centrifugation at 16,000 g for 30 minutes at 25 °C. The supernatant was collected and sodium chloride was added to a final concentration of 1 M. APOBEC3B-mycHis was allowed to bind to 50 µL nickel-NTA resin per 10x15cm plates for two hours at 4°C. The resin was collected in BioRad polyprep columns and washed with 25 mM HEPES pH 7.4, 300 mM sodium chloride, 0.1% Triton X-100, 40 mM imidazole, and 20% glycerol. Protein was eluted in the same buffer with the addition of TCEP to 1 mM and 300 mM imidazole. Purity and concentration were assessed by PAGE with Coomassie stain with gels imaged using a LI-COR Odyssey instrument.

**A3B activity assays.** Deamination reactions were performed at 37°C for 2 hrs using whole cell lysate, 4 pmol of oligonucleotide (5′-ATTATTATTATTCAAATGGATTTATTTATTTATTTATTTATTT-fluorescein), 0.025 U uracil DNA glycosylase (UDG), 1x UDG buffer (NEB), and 1.75 U RNase A. Reaction mixtures were treated with 100 mM NaOH at 95°C for 10 min to achieve complete backbone breakage. Reaction mixtures were separated on 15% Tris-borate-EDTA

(TBE)-urea gels to separate substrate from product. Gels were scanned using a Typhoon FLA-7000 image reader.

A3B activity assays with purified A3BmycHis or mutants were performed similarly as above in 25 mM HEPES pH 7.4, 50 mM NaCl, 0.4 U/mL Roche RNase Inhibitor for the indicated amounts of time at 37°C. Reactions were stopped at 95°C for 5 minutes then UDG was added to 0.4 U/reaction and incubated for 10 minutes at 37°C. Sodium hydroxide was added to 100 mM and reactions were heated to 95°C for 5 minutes. An equivalent volume of 80% formamide in 1x TBE with xylene cyanol and bromophenol blue was added and reactions were heated again to 95°C for 3 minutes to ensure melting of double stranded regions of DNA/RNA. Products were separated by 15% denaturing PAGE and digitally scanned using a LI-COR Odyssey imager. Quantitation was performed using LI-COR Odyssey software.

**Electrophoretic mobility shift assays.** For competition experiments, EMSAs were performed in 25 mM HEPES pH 7.4, 50 mM sodium chloride, 0.4 U/µL Roche RNase inhibitor. For R-loop substrate EMSAs NEB2 buffer (no BSA) was used to promote annealing of substrates. Oligonucleotide substrates (illustrated in **Fig. 7a** and full sequences listed in **Table S2**) were annealed by heating the components to 95°C in a heat block then permitted to cool to >10°C below the predicted annealing temperature under the buffer conditions (UNAFold). Reactions were set up with labeled oligo in the tube to which A3B or mutants were added to the appropriate concentration. Reactions were incubated at room temperature for 5 minutes and then either run or competitor was added with an additional 10 minutes incubation at room temperature. To run the gels, an equal volume of agarose gel loading dye (30% polyethylene glycol, 1x Tris-borate EDTA and dyes) was added to each reaction mix and half of each reaction was loaded on the gel. Gels were imaged using a LI-COR Odyssey and quantitated with LI-COR Odyssey software.

**Drug treatments.** PMA (Sigma Aldrich, P8139) was added to media at 25 ng/ml at 37°C with 5% $CO_2$ for denoted time. JQ1 (Tocris, 4499) was added to media at 0.5 μM at 37°C with 5%$CO_2$ for 4 hrs unless denoted otherwise. Triptolide (Tocris, 3253; Selleckchem, S3604) was added to media at 1 μM at 37°C with 5% $CO_2$ for 4 hrs unless denoted otherwise. Flavopiridol (Selleckchem, S1230) was added to media at 1 μM at 37°C with 5% $CO_2$ for 1 hr unless denoted otherwise. Doxycycline (MP Biomedicals, 198955) was added to media at 1 μg/mL at 37°C with 5% $CO_2$ for 24 hrs unless denoted otherwise. Pladienolide B (Tocris, 6070) was added to media at 5 μM at 37°C with 5% $CO_2$ for 2 hrs unless noted otherwise.

**Antibodies.** Primary antibodies used in these experiments were αTubulin (Sigma Aldrich, T5168 and Abcam, ab4074), αA3B (5210-87-13, in house[11]), αFlag (Sigma Aldrich, F1804), αTopoisomerase I (Abcam, ab109374), αLamin B1 (Abcam, ab16048), αIgG2a (Sigma Aldrich, M5409), αHA (Cell Signaling, 3724S), αGFP (Abcam, ab290, Lot# GR3251545 and GR3270983 for ChIP), αHNRNPUL1 (gift from Prof. Stuart Wilson, University of Sheffield, UK), αrabbit IgG Isotype Control (Invitrogen, 02-6102, lot#RI238244), αRNA/DNA Hybrid S9.6 (Kerafast, ENH001 or obtained in house from the hybridoma cell line), αdsDNA (Abcam, ab27156) and αgamma-H2AX

(Novus, NB100-384). Secondary antibodies used were αrabbit IRdye 800CW (LI-COR, 827-08365), αmouse IRdye 680LT (LI-COR, 925-68020), αrabbit HRP (Cell Signaling, 7074P2 or Sigma Aldrich, A0545), and αmouse HRP (Cell Signaling, 7076P2 or Sigma Aldrich, A8924), Alexa Fluor 488 goat anti-mouse IgG (Invitrogen, A-11029), Alexa Fluor 594 goat anti-mouse IgG (Invitrogen, A-11032), Alexa Fluor 488 goat anti-rabbit IgG (Invitrogen, A-11034) and Alexa Fluor 594 goat anti-rabbit IgG (Invitrogen, A-11037).

**Co-Immunoprecipitation.** Semi-confluent 293T cells were transfected with plasmids using TransIT-LT1 (Mirus) per manufacturer's protocol. Cells were harvested in 1xPBS 48 hrs post-transfection. Cells were washed two times in 1xPBS followed by lysis [150 mM NaCl, 50 mM Tris-HCl pH 8.0, 0.5% Tergitol, 1x Protease inhibitor (Roche), RNase and DNase]. Cells were vortexed vigorously and incubated at 4°C for 30 minutes prior to clearing by centrifugation. Cleared lysates were then added to anti-Flag M2 Magnetic Beads (Sigma M8823) followed by end-over-end rotation overnight at 4°C. Beads were then washed three times in lysis buffer followed by elution in elution buffer [lysis buffer + 0.15 mg/mL Flag Peptide (Sigma).

**EdU and PI Staining.** Semi-confluent MCF10A or U2OS cells were treated with 10uM EdU for 2 hours prior to harvesting. Click-iT Plus EdU Alexa Fluor 488 Flow Cytometry Assay Kit (Invitrogen C10632) with the addition of FxCycle PI/RNase Staining Solution (Invitrogen F10797) was used per manufacturer's protocol and flow cytometry was performed on BD LSRFortessa.

**RNA/DNA hybrid slot blots.** RNA/DNA hybrid slot blot experiments were performed as described[12,13]. RNase H sensitivity was carried out by incubation with 2 U of RNase H (NEB, M0297) per μg of genomic DNA for 18 hrs at 37°C. S9.6 and dsDNA samples were run on the same membrane and cut for primary antibody incubation. Images were acquired with LI-COR Odyssey Fc. Exposure setting for each antibody were consistent within experiments. S9.6 signal relative to dsDNA was quantified using Image Studio software (LI-COR Biosciences). Quantification was performed using S9.6 and dsDNA signal within in the linear range and normalized to wild-type, untreated or control samples.

**mRNA quantification.** Isolation of RNA and RT-qPCR methods and primers have been described[14]. The abundance of various mRNAs was quantified by RT-qPCR relative to the stable housekeeping transcript, *TBP*. Primer sequences are listed in **Table S2**.

**RNA/DNA immunoprecipitation (DRIP).** DNA/RNA immunoprecipitation was performed as described in refs.[15,16] using the S9.6 antibody[17]. Non-crosslinked nuclei were lysed in nuclear lysis buffer (50 mM Tris-HCl pH 8.0, 5 mM EDTA, 1% SDS) and subjected to Proteinase K treatment (Sigma Aldrich) for 3 hrs at 55ºC. Genomic nucleic acids were precipitated with isopropanol, washed in 75% ethanol and sonicated in IP dilution buffer (16.7 mM Tris-HCl pH 8.0, 1.2 mM EDTA, 167 mM NaCl, 0.01% SDS, 1.1% Triton X-100) with Diagenode Bioruptor to an average length of 500 bp. Following addition of protease inhibitors (0.5 mM PMSF, 0.8 μg/ml pepstatin A, 1 μg/ml leupeptin), sonicated genomic nucleic acids were precleared with protein A Dynabeads (Invitrogen) blocked with acetylated BSA (B8894, Sigma Aldrich). 10 µg were subjected to S9.6 or no antibody immunoprecipitation overnight at 4°C. RNase H sensitivity was carried out by incubation with 1.7 U RNase H (M0297; NEB) per µg of genomic DNA for 3 hrs at 37°C before IP. Retrieval of the immunocomplexes with beads, washes and elution were performed as described for ChIP. Samples were incubated with Proteinase K (Sigma Aldrich) at 45°C for 2 hrs. For qPCR analysis, DNA was purified with QIAquick PCR purification kit (QIAGEN) and analyzed by qPCR with Rotor-Gene® Q and QuantiTect SYBR green (QIAGEN). The amount of immunoprecipitated material at a particular gene region was calculated as the percentage of input after subtracting the background signal (no antibody control). The primers used for DRIP are listed in **Table S2**. For DRIP-Sequencing analysis, multiple S9.6 IPs were pooled. DNA was purified with MinElute® PCR purification Kit (QIAGEN) and subjected to library preparation and sequencing on a NovaSeq 6000 with 150 bp paired-end reads at Oxford Genomics Centre (WTCHG, University of Oxford).

**RNA/DNA hybrid and protein co-immunoprecipitation.** DNA/RNA hybrid co-IPs were carried out accordingly to ref.[16] using S9.6 antibody[17]. Non-crosslinked nuclei were lysed in RSB buffer (10 mM Tris-HCl pH 7.5, 200 mM NaCl, 2.5 mM MgCl2) with addition of 0.2% sodium deoxycholate, 0.1% SDS, 0.05% sodium lauroyl sarcosinate and 0.5% Triton X-100. Nuclear extracts were then sonicated with Diagenode Bioruptor and diluted in RSB with 0.5% Triton X-100 (RSB + T). RNA/DNA hybrids were immunoprecipitated for 2 hrs at 4ºC with BSA-blocked protein A Dynabeads (Invitrogen) conjugated with the S9.6 antibody in the presence of 1.2 ng of RNase A (PureLink, Invitrogen). Washes of the immunocomplexes were carried out with RSB + T (4 times) and RSB (2 times). Immunocomplexes were then eluted by incubating at 70°C with 1x LDS (Invitrogen) and 100 mM DTT for 10 min. Where indicated, IPs were performed in the presence of 1.3 μM DNA/RNA hybrid competitors, prepared as described[18] (ssDNA-CGGTGTGAATCAGAC; ssRNA-GUCUGAUUCACACCG). The same procedure was used for protein co-immunoprecipitation and anti-GFP antibody (ab290, Abcam) was used instead of S9.6 antibody. Proteins were separated by SDS-PAGE and immunoblotted with αA3B (5210-87-13, ref.[11]), αTopoisomerase I (Abcam, ab109374), αLamin B1 (Abcam, ab16048), αGFP (Abcam, ab290), and αHNRNPUL1 (gift from Prof. Stuart Wilson, University of Sheffield, UK) antibodies. For RNA/DNA hybrid slot blot analysis, A3B-eGFP co-IP was performed starting from 350 μg of proteins following the same procedure without the addition of RNase A. Immunocomplexes were eluted in 1% SDS and 0.1 M $NaHCO_3$ for 30 min at room temperature and nucleic acids were precipitated overnight with isopropanol and glycogen (Roche) after Proteinase K digestion (Sigma Aldrich) for 2 hrs at 45ºC. RNase H sensitivity was performed by incubating with 7.5 U of RNase H (NEB, M0297) for 2.5 hrs at 37°C.

**Chromatin immunoprecipitation (ChIP).** Chromatin immunoprecipitation experiments were carried out as described previously with few modifications[15,16]. Cells were crosslinked with 1% formaldehyde at 37ºC for 15 min before the reactions were quenched with 0.125 M glycine for 5 min. Nuclei were isolated by lysing cells with cell lysis buffer (5 mM PIPES pH 8.0, 85 mM KCl, 0.5% NP-40 supplemented with 0.5 mM

PMSF and 1X Complete EDTA-free protease inhibitors, Sigma Aldrich). Nuclear pellets were then resuspended in nuclear lysis buffer (50 mM Tris-HCl pH 8.0, 5 mM EDTA, 1% SDS supplemented with 0.5 mM PMSF and 1X Complete EDTA-free protease inhibitors, Sigma Aldrich) before sonication (Diagenode Bioruptor). Insoluble chromatin was removed by centrifugation. Soluble chromatin was then diluted in ChIP IP buffer (16.7 mM Tris-HCl pH 8.0, 1.2 mM EDTA pH 8.0, 167 mM NaCl, 0.01% SDS, 1.1% Triton X-100 supplemented with 0.5 mM PMSF and 1X Complete EDTA-free protease inhibitors, Sigma Aldrich) and precleared by incubation with protein A Dynabeads (Invitrogen) blocked with acetylated BSA (B8894, Sigma Aldrich). Precleared chromatin was then incubated with $\alpha$GFP antibody (ab290, Abcam). BSA-blocked protein A Dynabeads were then added to collect immunocomplexes and washed once with buffer A (20 mM Tris-HCl pH 8.0, 2 mM EDTA, 0.1% SDS, 1% Triton X-100 and 0.150 M NaCl), once with buffer B (20 mM Tris-HCl pH 8.0, 2 mM EDTA, 0.1% SDS, 1% Triton X-100 and 0.5 M NaCl), once with buffer C (10 mM Tris-HCl pH 8.0, 1 mM EDTA, 1% NP-40, 1% Sodium Deoxycholate and 0.25 M LiCl) and then twice with buffer D (10 mM Tris-HCl pH 8.0 and 1 mM EDTA). Chromatin complexes were eluted in 1% SDS and 0.1 M $NaHCO_3$. Samples were decrosslinked by incubating at 65°C for at least 4 hrs in the presence of RNase A (PureLink, Invitrogen) and NaCl (0.3 M) and digested with proteinase K (Sigma Aldrich) for 2 hrs at 45°C. DNA purification and qPCR analysis were performed as described for DRIP. For ChIP-sequencing analysis, multiple ChIP IPs were pooled. DNA was purified with MinElute® PCR purification Kit (QIAGEN) and subjected to library preparation and sequencing on a NovaSeq 6000 with 150 bp paired-end reads at Oxford Genomics Centre (WTCHG, University of Oxford).

**Immunofluorescence for R-loop analysis.** S9.6 immunofluorescence was performed as described[19]. Briefly, U2OS or MCF10A cells were treated with the transcription initiation inhibitor (triptolide, final concentration 1 μM) for 4 hrs or transfected with indicated constructs and treated with JQ1 (final concentration 0.5 μM) for 4 hrs or PladB (final concentration 5 μM) for 2 hrs. After incubation, cells were fixed with 100% ice-cold methanol at 4°C for 10 min and washed three times with PBS at room temperature. For *in vitro* RNase H treatment, fixed cells were washed with nuclease-free water to remove PBS and treated with 150 U/ml RNase H in 1X RNase H reaction buffer (NEB, #0297). Cells were incubated for 2 hrs at 37°C followed by 2X 5 min washes with 1X PBS. Untreated samples were similarly treated except using 1X RNase H reaction buffer without enzyme. Cells were then blocked with 3% BSA/TBS-T (0.05% Tween-20) at room temperature for 1 hr and incubated with S9.6 antibody (Kerafast, #ENH001, 1:200) at 4°C for 18 hrs. For detection of eGFP-overexpressed cells, cells were co-stained with GFP antibody (Abcam, ab290, 1:1000). Following primary antibody incubations, cells were washed with PBS 3X 5 min and incubated with appropriate secondary antibody in blocking buffer at room temperature. After 1 hr, cells were washed in PBS 3X 5 min and each cover slip was mounted on a 12 mm glass slide using vectashield mounting medium containing DAPI (Vector Labs, #H-1200). Samples were analyzed using a Fluoview 3000 confocal microscope (Olympus) or Nikon AR1 (University of Minnesota Imaging Center) and nucleoplasmic S9.6 signal was quantified by removing both cytoplasmic and nucleolar signal using Image J (v 1.48) as described in Quantification and Statistical Analysis section. All constructs were expressed to similar levels.

**Immunoblot analysis.** For immunoblotting assays, the samples were combined with 2.5x SDS-PAGE loading buffer. Samples were separated by a 4-20% gradient SDS-PAGE gel and transferred to PVDF-FL membranes (Millipore). Membranes were blocked in blocking solution (5% milk + PBS supplemented with 0.1% Tween20) and then incubated with primary antibody diluted in blocking solution. Secondary antibodies were diluted in blocking solution + 0.02% SDS. Membranes were imaged with a LI-COR Odyssey instrument or film.

**ChIP-seq and DRIP-seq data processing.** Adapters were trimmed with Cutadapt version 1.13[20] in paired-end mode with the following parameters: -q 15, 10 –minimum-length 10 -A AGATCGGAAGAGCGTCGTGTAGGGAAAGAGTGT -a AGATCGGAAGAGCACACGTCTGAACTCCAGTCA. Obtained sequences were mapped to the human hg38 reference genome with STAR version 2.6.1d[21] and the parameters --runThreadN 16 --readFilesCommand gunzip -c –k --alignIntronMax 1 --limitBAMsortRAM 20000000000 --outSAMtype BAM SortedByCoordinate. Properly paired and mapped reads (-f 3) were retained with SAMtools version 1.3.1[22]. PCR duplicates were removed with Picard MarkDuplicates tool. Reads mapping to the DAC Exclusion List Regions (accession: ENCSR636HFF) were removed with Bedtools version 2.29.2[23]. FPKM-normalized bigwig files were created with deepTools version 2.5.0.1[24] bamCoverage tool with the parameters –bs 10 –p max -e --normalizeUsing RPKM. ChIP-seq and DRIP-seq peaks were called with MACS2 version

2.1.1.20160309[25] and the parameters: callpeak -f BAMPE -g 2.9e9 -B -q 0.01 –call-summits. Each IP and its respective input were used as treatment and control, respectively. DRIP-seq differential peak calling was performed with MACS2 bdgdiff tool.

**Transcription unit annotation.** Gencode V31 annotation, based on the hg38 version of the human genome, was used to extract the location of the transcription units. All genes were taken from the most 5' transcription start site (TSS) to the most 3' poly(A) site / transcription end site (TES). The eRNAs annotation based on the hg38 version of the human genome was taken from the FANTOM5 database.

**Metagene profiles.** Metagene profiles were generated from FPKM-normalized bigwig files with Deeptools2 computeMatrix tool with a bin size of 10 bp and the plotting data obtained with plotProfile –outFileNameData tool. Graphs were then created with GraphPad Prism 8.3.1.

**RNA-seq data processing.** RNA-seq data from ref.[26] were processed as follow: adapters were trimmed with Cutadapt in single-end mode with the following parameters: -q 15, 10 –minimum-length 10 -a AGATCGGAAGAGCACACGTCTGAACTCCAGTCA –max-n 1. The trimmed reads were mapped to the human hg38 reference genome with STAR and the parameters --runThreadN 16 --readFilesCommand gunzip -c –k --limitBAMsortRAM 20000000000 --outSAMtype BAM SortedByCoordinate. SAMtools was used to retain only properly mapped reads (-F 4). Gene expression level (transcripts per millions, TPM) were calculated with Salmon version 0.13.1[27] and the

Gencode V31 annotation. For each gene, only the highest expressed transcript was retained.

**APOBEC mutation and gene expression.** Whole exome sequencing and RNAseq data sets for all primary breast tumor specimens (n = 977) and normal breast tissues (n = 111) in The Cancer Genome Atlas (TCGA) were downloaded from Broad Institute analysis pipeline through the Firehose GDAC resource (http://gdac.broadinstitute.org/). Similarly, whole genome sequencing data sets for all primary breast tumor samples (n = 794) in the International Cancer Genome Consortium (ICGC) were downloaded from the ICGC data portal (https://dcc.icgc.org/). Because ICGC tumors lack corresponding RNAseq data, expression values for genes in normal breast tissues were obtained by averaging available GTEx data (n = 29,589 genes from 396 normal breast tissue samples; https://gtexportal.org/home/).

Single base substitution (SBS) mutations from TCGA and ICGC breast cancers were used for analyses here (*i.e.*, INDELs and other more complex somatic variations were filtered out). Tumor data sets were ranked initially by APOBEC mutation enrichment scores using established methods[30]. Enrichment score significance was assessed using a Fisher's exact test with Benjamini-Hochberg false discovery rate (FDR) correction (q<0.05). TCGA breast tumors with significant APOBEC mutational signature enrichments (n = 154 tumors) were used to test whether mutation load per megabase associates with differential gene expression (tumor vs normal tissue). Mean normal expression values for each gene from 111 normal breast tissues from the TCGA breast cancer data set were used to generate a baseline for determining fold-changes in gene expression in tumor tissues. For each of the 154 APOBEC signature-enriched tumors, we first generated 7 gene expression groups: (1) tumor genes with expression values of 0; (2) tumor genes with expression values less than 0.8-fold of the normals (first quartile of all genes in all tumors); (3) tumor genes with fold changes between 0.8- and 1.2-fold of the normals (covers from first quartile to third quartile of all genes); (4) tumor genes with fold changes between 1.2- and 4-fold above the normals; (5) tumor genes with fold changes between 4- and 8-fold above the normals; (6) tumor genes with fold changes between 8- and 16-fold above the normals; (7) tumor genes with fold changes greater than 16-fold above the normals. Last, we calculated the fraction of APOBEC signature mutations (TCW to TTW or TGW) per tumor per megabase using the exon lengths of the genes in each group.

A similar analysis was done for ICGC tumor mutation vs GTEx expression values. Five expression groups were created: non-expressed genes in (Exp = 0) and all other genes divided into expression quartiles. Only C-to-G and C-to-T mutations in TCW trinucleotide motifs were used in these analyses, and were plotted for each expression group as 1) total number of T[C>G/T]W mutations, 2) total number of T[C>G/T]W mutations divided by the total number of all SBSs in a tumor, and 3) total number of T[C>G/T]W mutations as a fraction of the total nucleotide size of genes' (exons and introns) in that expression group (mutations per megabase per tumor). Gene size information was downloaded from the UCSC table browser resource (https://genome.ucsc.edu/cgi-bin/hgTables), and correspond to the "UCSC Genes, knownGene" reference set. All mutation calls and gene sizes/positions are relative to the hg19 human reference genome.

**Splice factor and APOBEC mutation analysis.** TCGA mutation data were downloaded from Broad GDAC Firehose as above. 119 splicing factor genes with recurring mutations in 33 cancers were used as the analysis gene set[31]. 107 of the 119 genes had deleterious mutations in the TCGA BRCA dataset. These deleterious mutations included stop codon mutations, splice site mutations, and insertion and deletion frameshift mutations. Trinucleotide contexts were calculated using the deconstructSigs package[32]. The APOBEC mutation signature in this analysis included all COSMIC SBS2 and/or SBS13 mutations[33]. Statistical analyses were done with Fisher's exact tests (with alpha = 0.05) and student's t-tests as indicated.

**APOBEC *kataegis* analysis using PCAWG WGS datasets.** To analyze APOBEC-associated kataegis, the set of whole-genome sequenced breast adenocarcinomas were downloaded from the official PCAWG release (https://dcc.icgc.org/releases/PCAWG; n = 198). *Kataegic* events were detected using a sample-dependent inter-mutational distance (IMD) cutoff, which is unlikely to occur by chance given the mutational burden and mutational pattern of each sample[29]. SigProfilerSimulator (v1.1.2) was used to generate a random distribution of the mutational spectra, while maintaining the +/-2bp sequence context and the strand coordination within genic regions of each mutation[28]. This background model was used to determine the cutoff for the sample-dependent IMD by ensuring that 90% of clustered mutations occur within the original sample compared to the expected distribution (q-value<0.01). The heterogeneity of mutation rates across the genome and the confounding effects of copy number alterations and clonality were addressed by performing a 10 mbp regional mutation density correction and by using a cutoff for the difference in variant allele frequencies between adjacent mutations in a clustered event (variant allele frequency difference <0.10)[29]. Clustered events consisting of ≥3 or ≥5 mutations were classified as *kataegis*. Events that did not fall within 10 kbp of a detected structural variant breakpoint were used for non-structural variation associated downstream analysis. All breakpoints were determined based on the official PCAWG release. Only base substitution mutations with TCW context were considered to be associated with APOBEC3 mutagenesis. A 1,000 bp window was included upstream and downstream of each DRIP-seq R-loop region to determine overlap with *kataegic* events. Mutation enrichment analysis was performed for each mutation by normalizing for the availability of a given motif (RTCA or YTCA) and the number of cytosines within +/-20 bp[30]. Additional analyses were conducted using R, Prism (v8.0), and the ggplot2 R package. Statistical significance between the tetranucleotide enrichments of *kataegis* and dispersed APOBEC3 mutation data sets was determined using a non-parametric Fisher's exact test, using an alpha of 0.05 (*P*-values reported in the text). Statistical significance for tetranucleotide mutation biases within samples containing overlaps of R-loop and *kataegic* events compared to dispersed mutations was assessed using a Mann-Whitney U-test (*Q*-values shown in each dot plot).

**Quantification and statistical analysis.** S9.6 immunofluorescence quantification was performed as described[19]. Briefly, nuclear S9.6 signals were quantified using Image J (v 1.48). First, the whole nucleus and nucleoli (overlaying DAPI-less regions, which correspond to nucleoli) S9.6 intensity values were measured, using ROI intensity by Image J. The nucleoli S9.6 intensity was excluded from the whole nucleus S9.6 intensity to obtain nuclear S9.6 values. For statistical analysis, one-way analysis of variance (ANOVA) was used when comparing more than two groups followed by a Dunnett multiple comparison test and for analyzing two groups, a Mann-Whitney test or two-tailed student's t-test was used as indicated. Statistical analyses for the bioinformatic studies are described as above.

**Data and code availability.** The accession number for the ChIP-seq, and DRIP-seq reported in this paper is GEO: GSE148581. The password for reviewers is: absdiyooplobjsj. The sequencing data generated by S9.6 IP experiments are available upon request to Reuben S. Harris.

**Table S1: Specific APOBEC3B interactors from AP-MS experiments**

See Microsoft Excel file titled “Supplementary Table 1” - Layer 1 for **Fig. 1a** A3B and S9.6 Interactors and Layer 2 for All Proteomics Hits.

**Table S2: Oligonucleotides**

| ***Primer Name*** | ***Sequence*** |
|---|---|
| A3B_shA3B-resistant-SDM-fwd | gaaaataaagaggggcagatcgaaccttctctgggat acaggggtctttcgaggcc |
| A3B_shA3B-resistant-SDM-Rev | ggcctcgaaagacccctgtatcccagagaaggttcgat ctgcccctctttattttc |
| A3B (full) Kozak, HindIII | nnnaagcttaccgccatgaatccacagatc |
| reverse a3bi KpnI | nnnnggtaccgtcgacgtttccctgattct |
| map4_fwd_EcoRI | nngaattcgggatggctgacctca |
| map4_rev_EcoRV | nngatatccggcttctgtagtatc |
| sf3b2_fwd_EcoRV | nngatatcccatggatgacccctc |
| sf3b2_rev_NotI | nngcggccgccgaaacttgaactc |
| hnrnpul1_fwd_EcoRI | nngaattcgggatggatgtgcgcc |
| hnrnpul1_rev_EcoRV | nngatatccctgtgtacttgtgcc |
| srsf7_fwd_KpnI | nnggtaccgggatgtcgcgttacg |
| srsf7_rev_EcoRV | nngatatccgtccattctttcagg |
| c1orf35_fwd_EcoRI | nngaattcgggatgttcggctcca |
| c1orf35_rev_EcoRV | nngatatccggcctcagagcctct |
| rbm8a_fwd_HindIII | nnaagcttcggatggcggacgtgc |
| rbm8a_rev_NotI | nngcggccgcccccggcgacgtct |
| gadd45a_qpcr37_fwd | agagcagaagaccgaaagga |
| gadd45a_qpcr37_rev | tgactcagggctttgctga |
| PHLDA1_qPCR24_fwd | cctccaactctgcctgaaag |
| PHLDA1_qPCR24_rev | aaatgtgctcgtcccacttc |
| HIST1H1E_qPCR60_fwd | gtcgggttccttcaaactca |
| HIST1H1E_qPCR60_rev | gctcctgctggcttcttg |
| DDX1_qPCR66_fwd | tgggaaagttacctacggtcag |
| DDX1_qPCR66_rev | aaatatccacatggcctttatagc |
| A3B_SDM_E255A_fwd | atgcggcactgcgcttctt |
| A3B_SDM_E255A_rev | aagaagcgcagtgccgcat |
| GADD45A_DRIP_fwd | tggactttcagccgagatgt |
| GADD45A_DRIP_rev | aggaattagtcacgggaggc |
| PHLDA1_DRIP_fwd | ggatggcctgacgattcttg |
| PHLDA1_DRIP_rev | atgaagaccgtggactgtgt |
| HIST1H1E_DRIP_fwd | cccaccgctctcagtaaaag |
| HIST1H1E_DRIP_rev | actcctctccccgactttgt |

| | |
|---|---|
| DDX1_DRIP_fwd | ggcctttccttccatcgga |
| DDX1_DRIP_rev | cacaaacgcaccggagaag |
| TFF1_DRIP_fwd | tatgaatcacttctgcagtgag |
| TFF1_DRIP_rev | gagcgttagataacatttgcc |
| Intergenic_DRIP_fwd | agtccataagcagccaaggt |
| Intergenic_DRIP_rev | aagacggggacagcattagt |
| JUNB_DRIP_fwd | ttaacagggaggggaagagg |
| JUNB_DRIP_rev | ttccacagtacggtgcagag |
| FOS_DRIP_fwd | ccatgacaggaggccga |
| FOS_DRIP_rev | acattcccaggaagagtacg |
| NAXE_DRIP_fwd | gcacaactctcgacctttgg |
| NAXE_DRIP_rev | ccacagatgaccaggacagt |
| ARL4D DRIP fwd | agagacggtgacccaag |
| ARL4D DRIP rev | cccatgctcctctctccttc |
| GAPDH_DRIP_fwd | ccactaggcgctcactgttc |
| GAPDH_DRIP_rev | tcgtagacgcggttcgg |
| Gemin7_DRIP_fwd | tcggtgagtacaaggtggtg |
| Gemin7_DRIP_rev | acctctttccgtcactcagg |
| DUSP1_DRIP_fwd | tgcccacttccatgaccat |
| DUSP1_DRIP_rev | ctctctcagtccaaaagcgg |
| HSPA8_DRIP_fwd | tgggtgaaagaaaacgctgg |
| HSPA8_DRIP_rev | acggaaatgtagggagtggg |
| HIST1H1B_DRIP_fwd | ttcgccttcttcggagtctt |
| HIST1H1B_DRIP_rev | aaactcaacaagaaggcggc |
| SYT8_DRIP_fwd | ccttgctcactcagtccaga |
| SYT8_DRIP_rev | gcctccttctcacctctgtt |
| PIM3_DRIP_fwd | aaatctgcttgtggacctgc |
| PIM3_DRIP_rev | aacagtcatcagcccctcc |
| A3B DNA EMSA | irdye800cwn-attagtcatatggat |
| A3B RNA EMSA | irdye800cwn-auuagucauauggau |
| DNA competitor | attagtcatatggat |
| RNA competitor | auuagucauauggau |
| A3B activity DNA | irdye700-gtgagagataggtggttgtatgatgat gaggtcaggatagtaggatagtgtggataggtgag |
| A3B activity complement | ctcacctatccacactatcctactatcctgacctcatcatc atacaaccacctatctctcac |

| | |
|---|---|
| A3B activity bubble | ctcacctatccacactatctgtaagttaagtaagttggtg<br>catacaaccacctatctctcac |
| A3B activity short R-loop | caccaacuuacuuaacuuaca |
| A3B activity long R-loop | aacaaccacgcacaccaacuuacuuaacuuaca |
| A3B activity RNA hybrid | cuacuauccugaccucaucau |

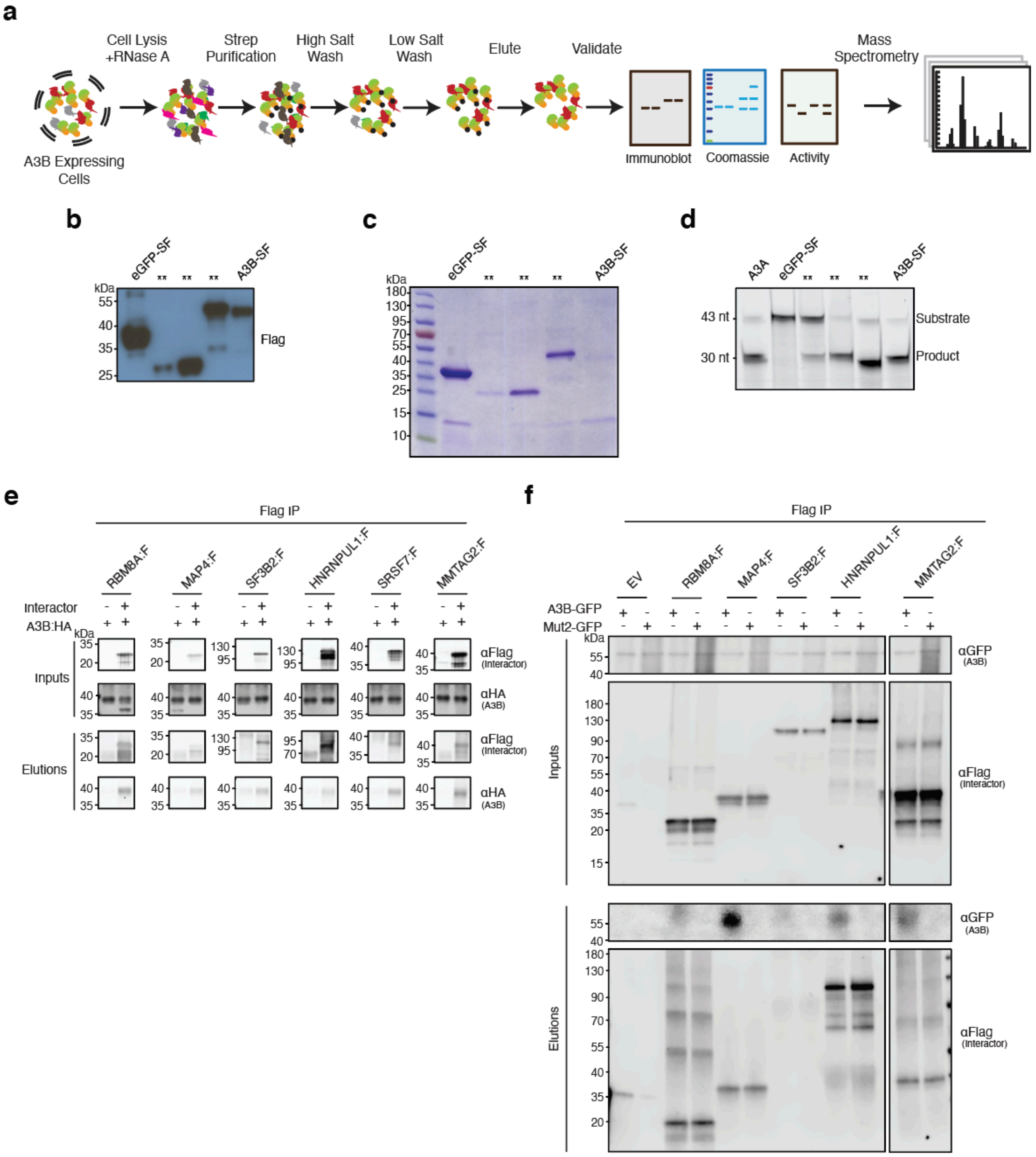


**Fig. S1. Controls for AP-MS experiments**

**a**, Schematic of the AP-MS workflow used to identfiy the cellular A3B interactome. A3B is shaded orange/green and cellular proteins are indicated by different shapes/colors.

**b-c**, Anti-Flag immunoblot and Coomassie gel analysis of eGFP-SF and A3B-SF following affinity purification and prior to analysis by mass spectrometry (**, samples that do not pertain to this manuscript, representative images).

**d**, DNA deaminase activity of eGFP-SF and A3B-SF following affinity purification (purified A3A was used as a positive control; **, samples that do not pertain to this manuscript, representative images).

**e**, co-IP of indicated Flag-tagged interactors and HA-tagged A3B in 293T cells. Upper immunoblots show the indicated proteins in whole cell lysates (input), and lower immunoblots show the Flag-immunoprecipitated samples (elution). kDa markers are shown the left of each blot and the primary antibody used for detection is shown to the right.

**f**, co-IP of indicated Flag-tagged interactors and eGFP-tagged A3B or eGFP-tagged Mut2 from 293T cells. Upper immunoblots show the indicated proteins in whole cell lysates (inputs), and lower immunoblots show the anti-Flag immunoprecipitated samples (elutions). kDa markers are shown to the left of each blot and the primary antibody used for detection is shown to the right.

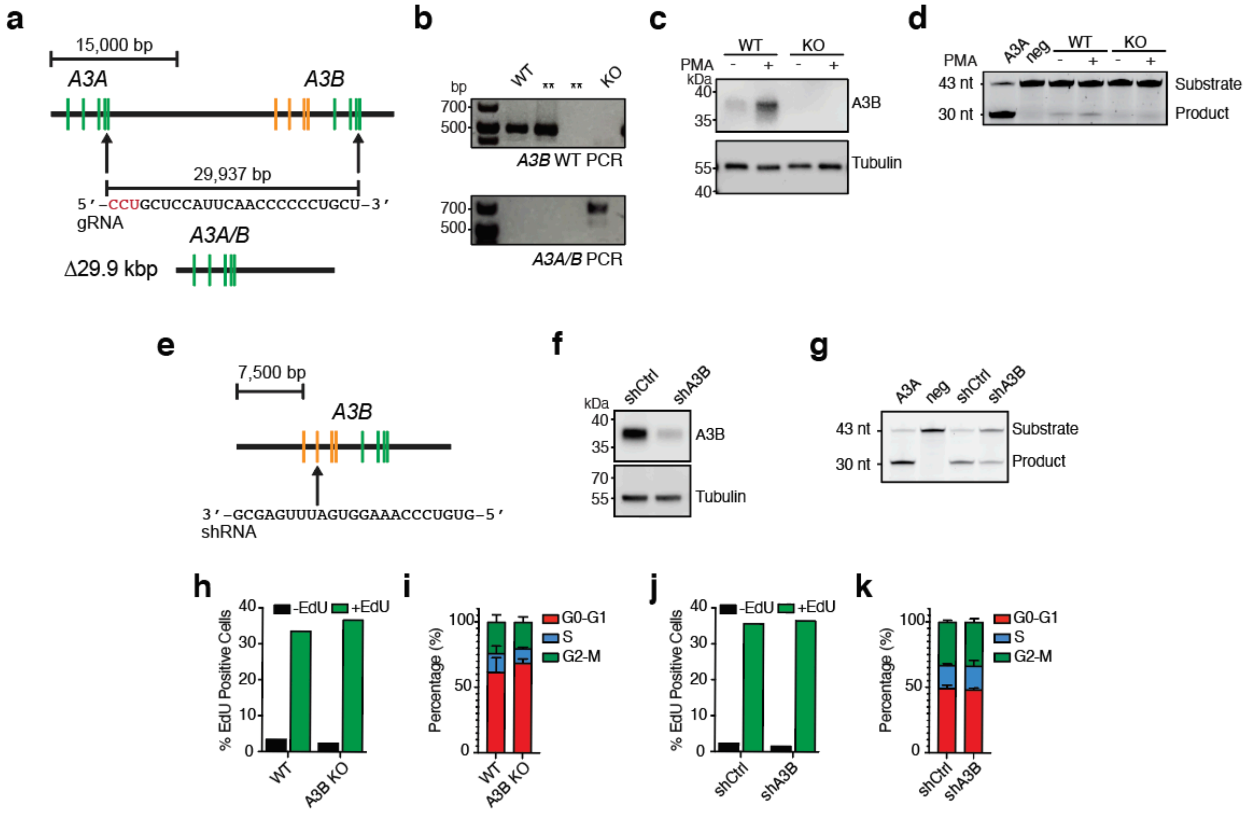


**Fig. S2. Cell line constructions and validations**

**a**, Schematic of the *A3B* knock-out strategy resulting in an *A3A*/*B* fusion gene (*i.e.*, coding sequence of *A3A* fused to 3'UTR of *A3B*). CRISPR cleavage sites are indicated by arrows and the gRNA-targeted region shared by the two genes is shown below with protospacer adjacent motif (red). Exons are indicated by colored boxes (N-terminal, orange; C-terminal, green).

**b**, Agarose gel analysis of PCR products that distinguish between wild-type *A3B* and 29.9 kbp *A3B* deletion allele. The wild-type (WT) *A3B* PCR uses one primer outside and one primer inside the 29.9 kbp deletion interval depicted in panel (a), whereas the knock-out (KO) PCR utilizes primers flanking the 29.9 kbp deletion (**, clones that do not pertain to this manuscript). *A3B* and *A3A/B* (KO) PCR products were confirmed by DNA sequencing.

**c**, Immunoblot of MCF10A wild-type (WT) and *A3B*-null derivative (KO) treated with DMSO or PMA (25 ng/ml, 24 hrs) and probed with the indicated antibodies.

**d**, DNA deaminase activity assay using extracts from MCF10A wild-type (WT) and *A3B*-null derivative (KO) treated with DMSO or PMA (25 ng/ml, 24 hrs; purified A3A was used as a positive control and reaction buffer as a negative control).

**e**, *A3B* gene schematic with an arrow indicating the exon 2 mRNA region targeted by an *A3B*-specific shRNA in depletion experiments (target sequence shown below). Exons are indicated by colored boxes (N-terminal, orange; C-terminal, green).

**f**, Immunoblot of U2OS shCtrl and shA3B cell lines probed with the indicated antibodies.

**g**, DNA deaminase activity assay of extracts from U2OS shCtrl and shA3B cell lines (purified A3A was used as a positive control and reaction buffer as a negative control).

**h**, EdU staining of MCF10A WT and A3B KO cell lines (n = 1 with a minimum of 10,000 cells per condition).

**i**, PI staining of MCF10A WT and A3B KO cell lines (n = 3 experiments with a minimum of 10,000 cells per condition; mean ± SD).

**j**, EdU staining of U2OS shCtrl and shA3B cell lines (n = 1 with a minimum of 10,000 cells per condition).

**k**, PI staining of U2OS shCtrl and shA3B cell lines (n = 3 experiments with a minimum of 10,000 cells per condition; mean ± SD).

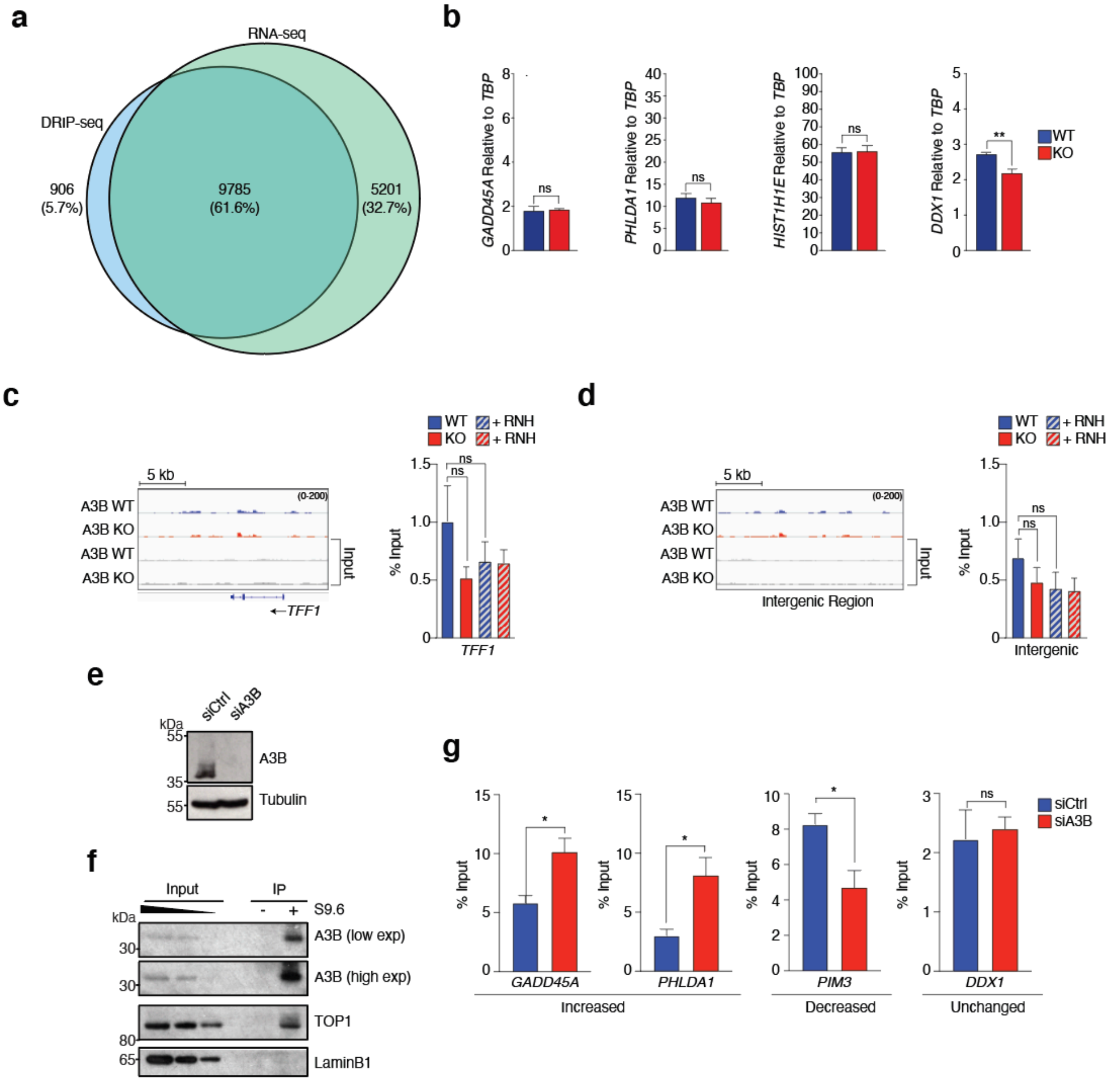


**Fig. S3. Supporting data for DRIP-seq experiments**

**a**, Venn diagram depicting the overlap between DRIP-seq positive genes and expressed genes (RNA-seq) in MCF10A (a similar overlap was observed for data sets from *A3B*-null cells, not shown).

**b**, RT-qPCR analysis of mRNA levels in MCF10A (WT) and *A3B*-null MCF10A (KO) cells. Values for the indicated genes are expressed relative to the housekeeping gene, *TBP* (n = 3; mean ± SEM; **, $P < 0.01$ by two-tailed unpaired t-test; ns, not significant).

**c-d**, DRIP-seq profiles for an untranscribed gene, *TFF1*, and an intergenic region in MCF10A (WT and *A3B*-null) cells. Independent quantification by DRIP-qPCR -/+ RNase H (RNH) treatment (striped bars) is shown in histograms to the right (n ≥ 4; means ± SEM expressed as percentage of input; ns, not significant).

**e**, Immunoblot of HeLa cells transfected with either an siRNA against Luciferase (siCtrl) or A3B (siA3B) and probed with the indicated antibodies.

**f**, S9.6 IP from HeLa cells. Inputs are shown on the left and the S9.6-immunoprecipitated samples (elution) on the right. kDa markers are shown on the left of each blot and the primary antibody used for detection is shown to the right. Lamin B1 is a negative control.

**g**, Independent quantification of HeLa S9.6 IP by DRIP-qPCR of genes from the subgroups listed in Fig. 5c-e. ($n \geq 3$; means ± SEM expressed as percentage of input; *, $P < 0.05$ by two-tailed unpaired t-test, ns, not significant).

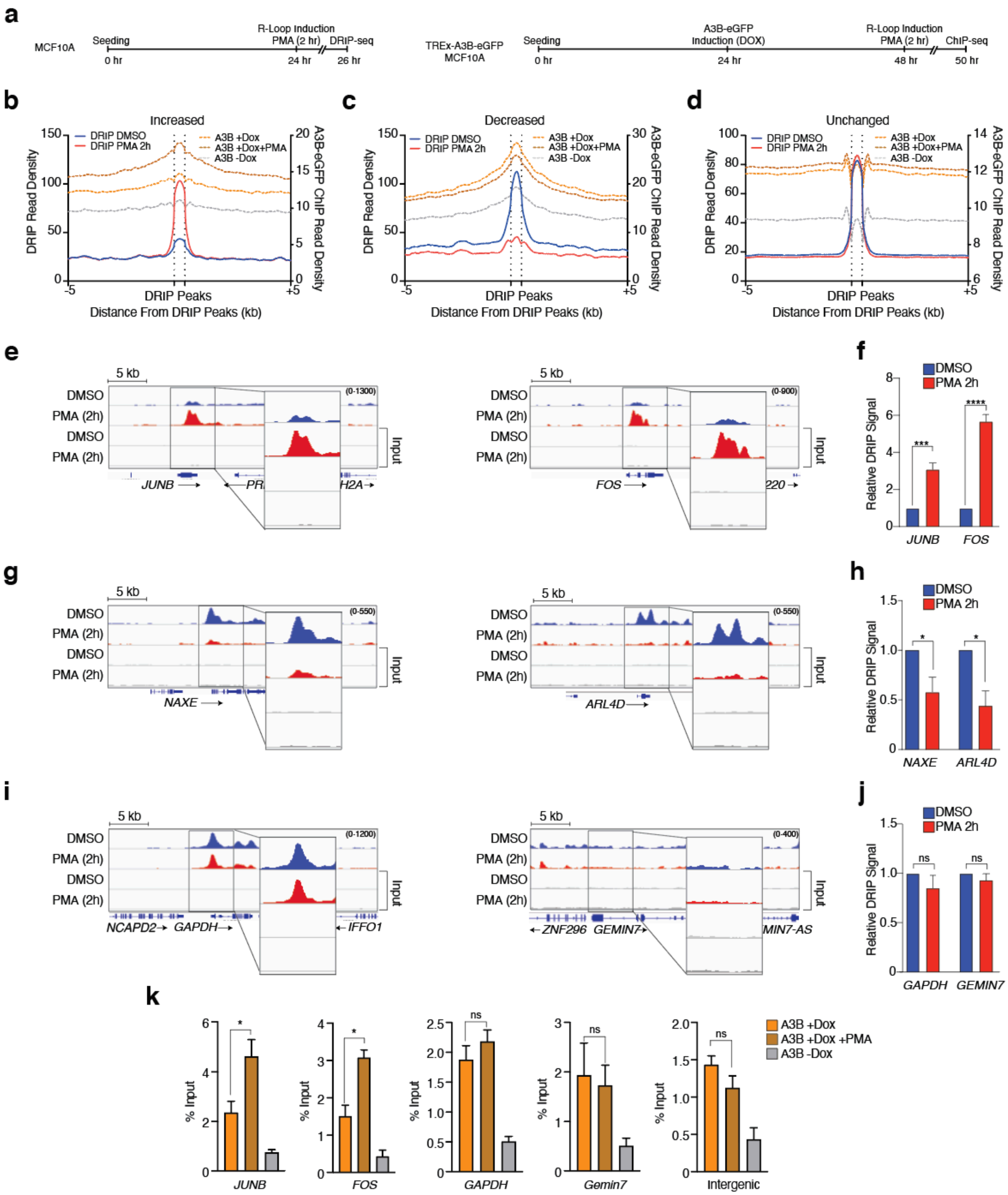


### Fig. S4. Kinetics of R-loop induction and resolution

**a**, Schematic of the DRIP-seq and A3B-eGFP ChIP-seq workflow used for panels b-j.

**b-d**, Meta-analysis of read density (FPKM) for DRIP-seq results from DMSO (blue) or PMA-treated MCF10A (red) partitioned into 3 groups (increased, decreased, and unchanged) as described in the text. A3B-eGFP ChIP-seq data (Dox-, Dox+, and

Dox+PMA in grey, orange, and brown dashed lines, respectively) superimposed on DRIP peaks -/+ 5 kb (right y-axis).

**e-f**, DRIP-seq profiles for *JUNB* and *FOS* from the increased data set in panel b. Independent quantification by DRIP-qPCR is shown in the histogram to the right (n = 4; means ± SEM normalized to DMSO; ***, $P < 0.001$, ****, $P < 0.0001$ by two-tailed unpaired t-test).

**g-h**, DRIP-seq profiles for *NAXE* and *ARL4D* from the decreased data set in panel c. Independent quantification by DRIP-qPCR is shown in the histogram to the right (n = 4; means ± SEM normalized to DMSO; *, $P < 0.05$ by two-tailed unpaired t-test).

**i-j**, DRIP-seq profiles for *GAPDH* and *GEMIN7* from the unchanged data set in panel d. Independent quantification by DRIP-qPCR is shown in the histogram to the right (n ≥ 3; means ± SEM normalized to DMSO; ns, not significant by two-tailed unpaired t-test).

**k**, Independent quantification by ChIP-qPCR is shown in the histogram for PMA-responsive (*JUNB, FOS*) and PMA non-responsive (*GAPDH* and *GEMIN7*) genes as well as an intergenic control (n ≥ 2; means ± SEM expressed as percentage of input; *, $P < 0.05$, ns, not significant by two-tailed unpaired t-test).

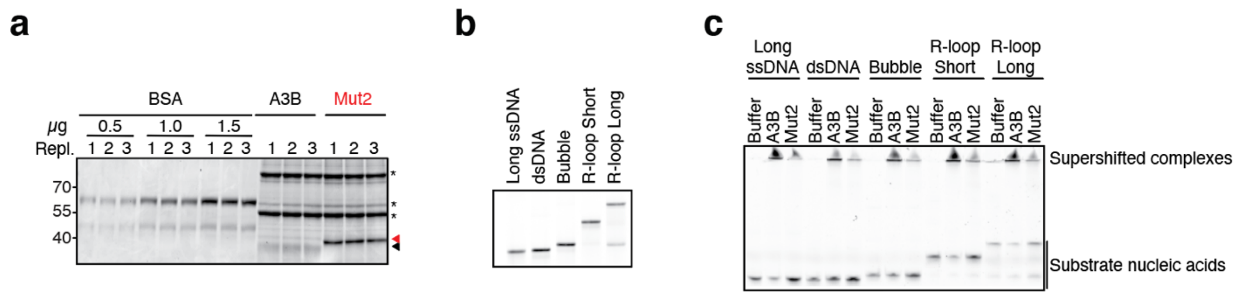


**Fig. S5. Purification and EMSAs for A3B and Mut2**

**a**, Coomassie-stained gel of Ni-NTA affinity purified A3B and Mut2 proteins from 293T cells (1, 2 and 3 are replicate loadings of the indicated proteins for purposes of quantification). kDa size standards are indicated on the left. Black and red arrow heads on the right indicate wildtype A3B-mycHis and Mut2-mycHis proteins, respectively (*, shared copurifying proteins).

**b**, Native TBE-PAGE of the 5' fluorescently labeled substrates depicted in Fig. 7a (size standards not applicable due to native conditions and different nucleic acid structures).

**c**, Native EMSA comparing wildtype A3B and Mut2 binding to the indicated nucleic acid substrates (*i.e.*, ssDNA, dsDNA, bubble, short R-loop, and long R-loop). In all instances, wildtype A3B binds more strongly than Mut2 indicated by more supershifted substrates, more intense staining of complexes retained in the wells, and larger diffusion "tails" within each well (an unavoidable issue if some of the complexes fail to enter the native acrylamide gel matrix). As in panel b, size standards are not applicable due to native conditions and different nucleic acid structures.

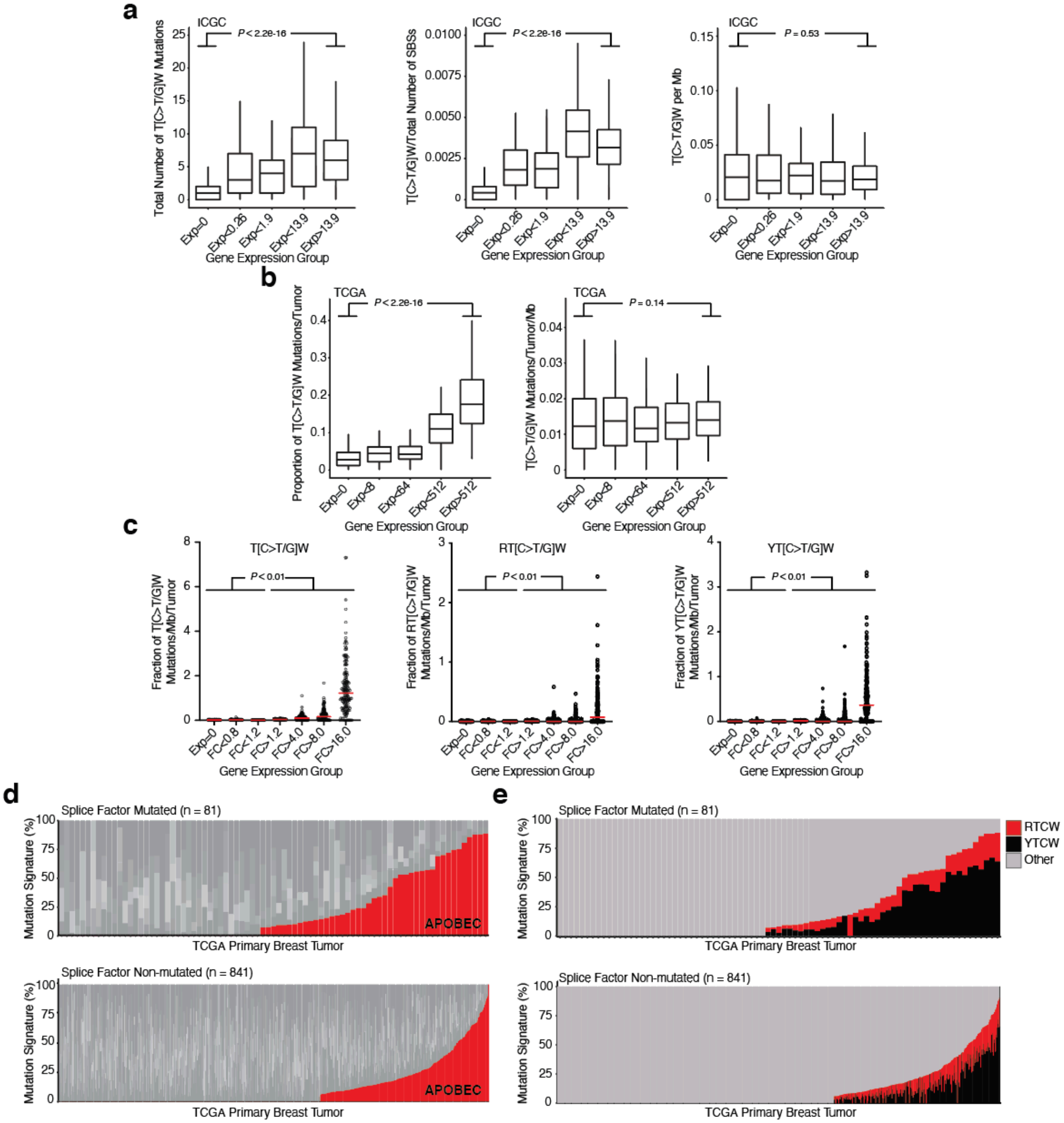


**Fig. S6. Additional analyses supporting model for R-loop mutation by A3B**

**a**, Positive correlations between gene expression levels and APOBEC signature T(C>T/G)W mutation number (left; $P < 2.2 \times 10^{-16}$) and frequency (middle; $P < 2.2 \times 10^{-16}$) in breast cancer ICGC data sets flatten upon normalization for gene size (right; $P = 0.53$; Pearson's correlation). Expression groups are based on gene expression levels in normal breast tissue from the Genotype-Tissue Expression (GTEx) project.

**b**, A positive correlation between gene expression levels and APOBEC signature T(C>T/G)W mutation number (left; $P < 2.2 \times 10^{-16}$) in TCGA breast cancer data sets flatten upon normalization for gene size (right; $P = 0.14$; Pearson's correlation).

Expression groups are 0 and quartiles for anything >0 and based on average expression levels for each gene using TCGA RNA-seq values from primary breast tumors.

**c**, Dot plot representations of the relationship between APOBEC signature mutations (per mb per tumor) and the indicated TCGA breast cancer gene expression groups (FC, fold-change relative to mean normal expression value in the TCGA normal breast tissue RNA-seq data). Left is identical to main **Fig. 8b** showing all APOBEC signature SBS, and the center and right panels show the RTCW and YTCW subsets of mutations, respectively. For each analysis, the pairwise comparisons are significant for all combinations of the lowest 4 and the highest 3 FC expression groups ($P \leq 3.5 \times 10^{-4}$ by Welsh's t-test).

**d**, Stacked bar graphs showing the proportion of each COSMIC mutation signature in TCGA breast tumors with mutations in splice factor genes or not (n = 81 and n = 841, respectively). Splice factor mutant tumors are more likely to have an APOBEC3 signature than controls (43/81 versus 326/841; *P* = 0.017, Fisher's exact test). COSMIC signatures SBS2 and SBS13 were combined for the APOBEC3 signature percentage (red), and other mutation signatures are shown in different shades of gray. The data here are identical those in **Fig. 8c** to facilitate comparison with tetranucleotide breakdowns in panel e.

**e**, An alternative representation of the data in panel d, with RTCW mutation proportions shown in red and YTCW mutation proportions in black. This analysis revealed a significant trend with only 1/43 (2.3%) of the APOBEC3 signature-enriched splice factor mutant breast tumors lacking mutations in A3B-associated RTCW motifs in comparison to 52/326 (15.9%) of the APOBEC3 signature-enriched non-splice factor mutant tumors (*i.e.*, the A3B-associated tetranucleotide preference is enriched in the splice factor mutant group and/or depleted from the non-splice factor mutant group; *P* = 0.028 by Fisher's exact test).
